# Mapping Parkinson’s Disease Progression through Deep Plasma Proteomics and Functional Genomics

**DOI:** 10.64898/2026.01.19.699565

**Authors:** Ji-sun Kim, Chanhee Jeong, Maria Seo-Hyun Kim, Hyeyeon Kim, Seungmin Lee, Kyeong Beom Jo, Wen Luo, Irina Shlaifer, Su Hyeon Ha, Saba Sane, Kyung Ah Woo, Emma Lee, Youngjin Lee, SooYeon Chae, Jisu Han, Atina G. Cote, JongHyun Seo, Gunwoo Park, Myungjin Lee, Gyurie Lee, Nidhi Sahni, Jung-Hyun Choi, Ji-Hwan Park, David E. Hill, Hunsang Lee, Kenneth A. Matreyek, Stefanie N. Kairs, Blake L. Tsu, Sangtae Kim, Christopher A. Barnes, Jean-François Trempe, Janusz Rak, Jung Hwan Shin, Thomas M. Durcan, Ki-Jun Yoon, Dae-Kyum Kim, Han-Joon Kim

## Abstract

Parkinson’s disease (PD) is a progressive neurodegenerative disorder lacking disease-modifying therapies, and its clinical management is limited by the absence of accessible biomarkers for tracking disease progression and treatment response. To map these complex disease trajectories, we implemented an ultra-deep plasma proteomics workflow integrating Mag-Net extracellular vesicle enrichment with Orbitrap Astral mass spectrometry to profile longitudinal samples from PD patients. This approach quantified 6,481 plasma proteins at unprecedented depth in PD studies, revealing distinct signatures directly associated with disease duration and dopaminergic therapy exposure. Candidate biomarkers were subsequently validated in an independent cohort using ELISA, demonstrating robust predictive utility in AI-driven prediction models. To uncover the mechanistic drivers underlying these systemic changes, we intersected our proteomic data with novel proteome-wide gene overexpression perturbation screens designed to identify regulators of alpha-synuclein pre-formed fibril (PFF) uptake and PFF-induced neuronal toxicity. Finally, an integrative network analysis combining three independent proteome-wide assays revealed that key pathological hubs, such as CD14, IFNG, and PLAT, are targets of currently approved pharmacological agents. Collectively, these findings provide a comprehensive, systems-level map for PD biomarker discovery and highlight druggable pathways to advance precision medicine strategies.

## Introduction

Parkinson’s disease (PD) is a progressive neurodegenerative disorder affecting over 8.5 million individuals globally, a prevalence that continues to rise^1,2^. While clinically defined by hallmark motor symptoms, PD is also characterized by pervasive non-motor manifestations, including cognitive impairment, sleep disturbances, depression, and autonomic dysfunction, which significantly impair quality of life^2–5^. These motor deficits manifest only after substantial dopaminergic neuronal loss, which marks the culmination of a prolonged prodromal phase that precedes diagnosis by years^6,7^. Current treatment options, such as Levodopa (L-DOPA), provide symptomatic relief by restoring dopamine levels^8–10^ but fail to halt or reverse disease progression^11^. Furthermore, clinical monitoring relies heavily on the Unified PD Rating Scale (UPDRS) and Hoehn and Yahr (H-Y) scale; these tools suffer from inherent subjectivity and inter-rater variability, and they inadequately capture non-motor symptoms or underlying molecular heterogeneity^12^. Consequently, there is an urgent need for robust biomarkers capable of enabling early diagnosis, tracking disease trajectory, and guiding therapeutic development.

Cerebrospinal fluid (CSF) offers a direct window into pathological processes within the central nervous system (CNS) and is considered an optimal source for biomarkers; however, its invasive collection precludes routine use in clinical monitoring^13,14^. In contrast, plasma is accessible via a simple blood draw and serves as a dynamic repository of proteins secreted or shed from tissues, offering systems-level insights^15^. Plasma proteomics thus holds promise for scalable screening and longitudinal monitoring in clinical trials^16,17^. Recent targeted mass spectrometry assays have successfully identified plasma panels, including Granulin precursor and Complement C3, that distinguish PD patients from prodromal controls^18^, while large-scale population studies have detected protein signatures predictive of PD decades before diagnosis^19^.

Despite these advances, two major hurdles impede the clinical translation of plasma proteomics in PD. First, the blood-brain barrier (BBB) acts as a selective gatekeeper that severely limits the release of brain-derived molecules into the peripheral circulation^20^. Consequently, CNS-specific biomarkers are often diluted to sub-detectable concentrations, obscured by the overwhelming dominance of high-abundance plasma proteins such as albumin and immunoglobulins^21^. Historically, this extreme dynamic range issue has restricted the analytical depth of plasma proteomics to the top 300-3,000 proteins, effectively masking the low-abundance signals essential for monitoring neurodegeneration^22^. Second, even when deep profiling identifies differential protein abundances, the resulting candidate lists often contain hundreds of targets^23^. Distinguishing true pathogenic drivers from downstream “bystanders”, proteins that change merely due to secondary cellular stress or inflammation, remains a significant bottleneck^24^. Without functional validation, these correlative markers often fail to translate into effective therapeutic targets.

To overcome these challenges, we integrated the Mag-Net membrane-bound particle enrichment method^25^ with ultra-high sensitivity Orbitrap Astral mass spectrometry^26,27^. This workflow allowed for deep longitudinal profiling of 20 plasma samples from 10 PD patients (**Supplementary Table S1**), quantifying 6,481 proteins across diagnostic and progressive stages. We leveraged these data to build predictive machine learning models for disease progression and, crucially, to bridge the gap between correlation and causation by integrating our proteomics with proteome-wide overexpression screens^28^. These screens identified novel regulators of alpha-synuclein pre-formed fibril (PFF) uptake and cytotoxicity^29,30^, while a meta-analysis of public RNA-sequencing datasets^31^ isolated candidates responsive to L-DOPA therapy. By synergizing deep proteomic profiling with functional genomics, the integrated network reveals novel, actionable biomarkers and elucidates the mechanisms driving PD progression and treatment response, establishing a foundation for precision therapeutic interventions.

## Results

### Deep plasma proteomics enables differentiation of longitudinal PD cohorts

We employed an integrated proteomics workflow combining Orbitrap Astral mass spectrometry^26,27^ with Mag-Net^25^, a technique designed to deplete abundant plasma proteins while enriching extracellular vesicles (EVs). EVs have emerged as attractive candidates for neurodegenerative disease biomarker discovery because they are actively secreted by neurons and glial cells^32,33^, traverse the blood–brain barrier^34,35^, and retain molecular signatures of central nervous system pathology in peripheral biofluids^36–38^. This platform enabled the quantification of 6,481 plasma proteins (**Fig. 1A; Supplementary Table S2**), representing the most comprehensive proteomic coverage of PD reported to date^39–42^. Dimensionality reduction techniques, including principal component analysis (PCA; **Fig. 1B**), t-distributed stochastic neighbor embedding (t-SNE; **Supplementary Fig. S1A**), and uniform manifold approximation and projection (UMAP; **Supplementary Fig. S1B**), revealed distinct clustering of plasma samples collected at diagnosis versus those collected after disease progression. These findings support the existence of stage-specific proteomic signatures that reflect underlying pathophysiological changes.

**Figure 1.**
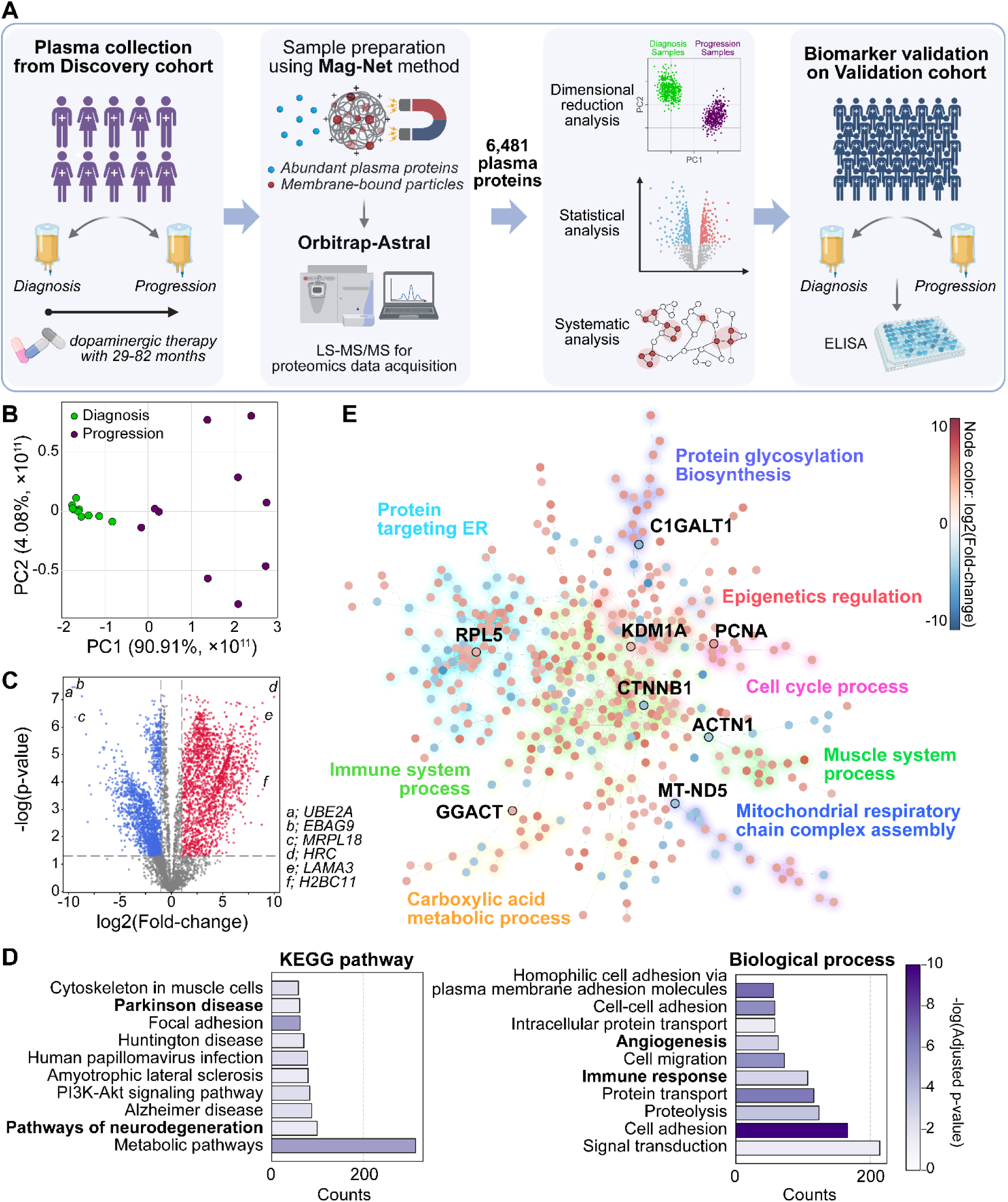
Longitudinal analysis of the plasma proteome of PD patients. **A.** Schematic overview of the study design, created with BioRender. **B.** PCA plot showing the distinct clustering of diagnosis and progression groups. **C.** Volcano plot showing DEPs identified by comparison of the diagnosis and progression group, with the thresholds of |log_2_FC|>1 and *p*<0.05. Red represents upregulated, and blue represents downregulated plasma proteins in the progression. The top 3 upregulated/downregulated proteins were annotated. P-value was calculated by an empirical t-test. **D.** Functional enrichment analysis of DEPs. The top 10 KEGG pathways and GO Biological process terms were identified. **E.** A SAFE analysis identifies 8 clusters within the network that represent specific GO biological process terms based on the similarity of their enrichment landscapes. Node colors indicate log2 fold changes.

To evaluate whether our workflow improved cohort resolution compared to conventional methods, we re-analyzed published longitudinal plasma and CSF proteomics data from the Parkinson’s Progression Markers Initiative (PPMI)^43^ using our analysis pipeline (**Materials and Methods**). These external datasets yielded ∼300 proteins per sample, a depth insufficient to separate diagnostic from progressive stages (**Supplementary Fig. S1C**). A power analysis confirmed that our cohort size provided robust statistical power (>0.8) for detecting differences between the diagnostic and progression groups^44^. In contrast, the achieved power values estimated from the PPMI CSF and plasma datasets were 0.074 and 0.088, respectively. These results suggest that our deep plasma proteomics pipeline may enable adequately powered analyses with smaller cohort sizes compared with conventional PPMI CSF or plasma proteomics datasets^45^. This contrast underscores the necessity of deep proteomic profiling to capture biologically meaningful differences in PD progression.

Next, we identified differentially expressed proteins (DEPs) between the diagnosis and progression samples. We observed high intra-group correlation and consistent inter-group divergence, confirming the robustness of our measurements (**Supplementary Fig. S1D**). Notably, samples from patients 03, 05, and 08, who had shorter disease durations (29, 36, and 45 months, respectively), exhibited stronger correlations with their respective baseline samples compared to patients with longer disease durations (>60 months). This indicates that the magnitude of plasma proteomic remodeling is significantly influenced by disease duration (**Supplementary Table S1**). A volcano plot of DEPs identified H2BC11 as a top upregulated protein; this nucleosomal histone is implicated in the epigenetic regulation of PD-associated genes, including Parkin, SNCA, and LRRK2^46^ (**Fig. 1C; Supplementary Table S2**). Conversely, UBE2A, a ubiquitin-conjugating enzyme linked to impaired Parkin-dependent mitophagy^47^, was among the most downregulated targets. Interestingly, alpha-synuclein (SNCA) was downregulated in progression samples [log_2_Fold-Change (log_2_FC) = −0.689 with p=0.0004]. Although blood RNA levels of alpha-synuclein have been reported to increase with disease duration^48^, plasma protein levels were shown to slightly decrease from H-Y stage I to stage II in patients with PD^49^, which is consistent with our findings.

To explore the biological relevance of these DEPs, we performed Gene Ontology (GO) and KEGG pathway enrichment analyses^50^ (**Fig. 1D; Supplementary Fig. S1E**). Enrichment of EV-associated terms validated the capacity of Mag-Net to selectively isolate vesicular proteins^25^. Additional enrichment of ER, Golgi, and mitochondrial components aligned with known mechanisms of alpha-synuclein toxicity, including ER–Golgi trafficking disruption, oxidative stress, and mitochondrial impairment^51^. Molecular function terms such as calcium ion binding and ATP binding further supported mitochondrial involvement^52^ (**Supplementary Fig. S1E**). Enriched biological processes included immune response^53^, angiogenesis^54^, protein transport^55^, and cell-cell communication^56^, all of which are implicated in PD pathogenesis (**Fig. 1D**).

To delineate functional protein networks, we applied Spatial Analysis for Functional Enrichment (SAFE)^57^ using a stringent threshold of |log_2_FC|>4 and p<0.05 (**Fig. 1E**). This analysis revealed eight core clusters enriched for proteins involved in: (i) protein glycosylation, (ii) epigenetic regulation, (iii) cell cycle, (iv) muscle system, (v) mitochondrial function, (vi) carboxylic acid metabolism, (vii) immune system, and (viii) ER biology. Key hub proteins with high betweenness centrality included RPL5, C1GALT1, PCNA, KDM1A, CTNNB1, ACTN1, MT-ND5, and GGACT, many of which have established roles in PD or other neurodegenerative disorders^58–65^. Collectively, these findings demonstrate that our high-resolution plasma proteomics platform not only enables reproducible differentiation of longitudinal PD cohorts (**Fig. 1B; Supplementary Fig. S1A-C**), but also uncovers biologically relevant pathways and candidate biomarkers, providing a foundation for mechanistic insights (**Fig. 1D; Supplementary Fig. S1E**).

### Multivariate analysis reveals disease progression and dopaminergic treatment as key determinants of the plasma proteome in PD

Previous plasma proteomics studies in PD primarily compared healthy controls to patients, focusing on binary disease status markers^18,19,66–70^. In contrast, our longitudinal design accounts for confounding variables. It reveals dynamic proteomic changes influenced by disease duration (ranging from 2 to 7 years), demographic factors, and evolving medication regimens (**Fig. 2A**). Furthermore, a PCA of fold changes across our longitudinal cohorts showed that the first principal component (**PC1 in Fig. 2A**) explained only about one-third of the total variance (**Supplementary Fig. S2A**). This indicates that longitudinal protein alterations are shaped by diverse biological and clinical factors, extending far beyond a simple binary distinction of disease status.

**Figure 2.**
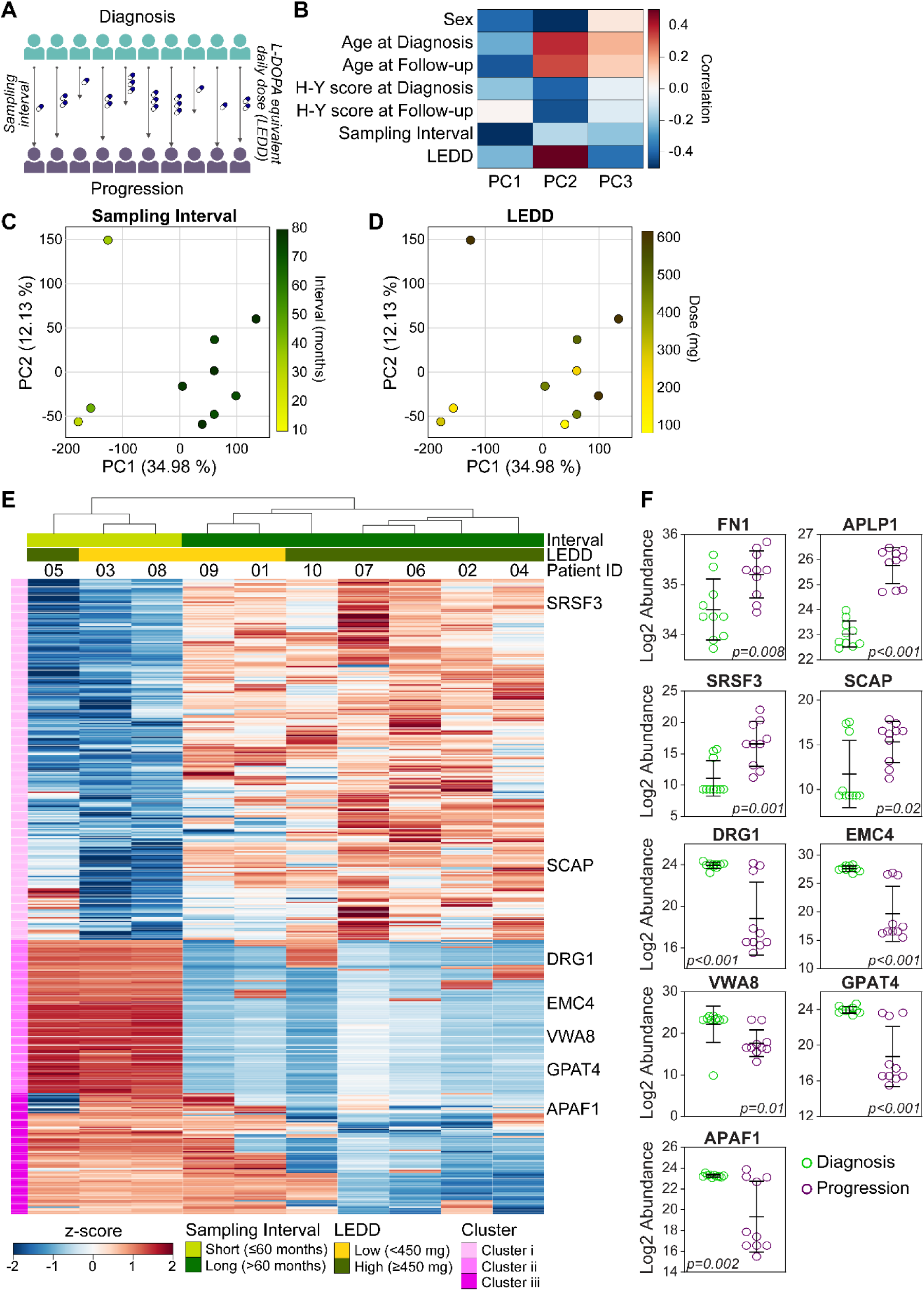
Impact of disease duration and medication on longitudinal plasma proteomic remodeling. **A.** Schematic illustration of calculating FC for each patient, created with BioRender. **B.** Heatmap of correlation between 7 covariates (sex, age at diagnosis, age at follow-up, sampling interval, and LEDD) and PCs. **C & D.** PCA plots of FCs for individual patients. Coloring represents (**C**) sampling interval and (**D**) LEDD level corresponding to the sample source. **E.** Hierarchical cluster map of the top 415 DEPs most discriminative for sampling interval or LEDD. The top 300 most discriminative DEPs for the sampling interval and all 123 discriminative DEPs for LEDD were picked, and the overlapped ones were excluded. The horizontal axis denotes patient IDs, and the vertical axis shows clusters of DEPs. **F.** Boxplot of protein abundance for 9 candidate biomarkers in discovery cohorts. The mean values and standard deviations in each group are shown as black lines. An empirical t-test was used to analyze the differences between groups for each candidate.

To further explore these influences, we conducted covariate analysis using patient metadata, including i) sex, ii) age at diagnosis/follow-up, iii) H-Y score, iv) sampling interval, and v) L-DOPA equivalent daily dose (LEDD)^71^ (**Supplementary Table S1**). Among the seven variables assessed, sampling interval and LEDD emerged as significant contributors to PC1 and PC2, respectively (**Fig. 2B**). Stratifying PCA plots by sampling interval revealed distinct separation along PC1, with shorter intervals associated with unique proteomic profiles (**Fig. 2C**). Additionally, PC2 values showed a correlation with increasing LEDD (**Fig. 2D**).

We subsequently identified 415 DEPs that were most discriminative for either sampling interval or LEDD (**Fig. 2E**). Hierarchical clustering highlighted three distinct groups: proteins (i) downregulated or (ii) upregulated in patients with shorter disease progression, and (iii) proteins modulated by LEDD. Notably, in clusters (i) and (ii), DEPs from patients with shorter progression intervals clustered tightly together, reflecting a coherent pattern of regulation. In contrast, DEPs from patients with longer progression intervals displayed more heterogeneous expression patterns, suggesting greater inter-individual variability at later disease stages. LEDD also significantly influenced clustering; patients receiving higher daily doses exhibited a distinct expression signature that aligned with cluster (iii). Together, these patterns indicate that both the disease-progression interval and LEDD leave detectable, distinct footprints in the plasma proteome.

To evaluate whether other datasets support similar stratification, we analyzed public longitudinal proteomics data from the PPMI^43^. While covering multiple time points (24 to 84 months), fold-change comparisons between baseline and follow-up samples in the PPMI cohort did not exhibit consistent temporal patterns in either CSF or plasma (**Supplementary Fig. S2B**). This observation may be attributable to differences in proteomic depth; the PPMI dataset detected approximately 500 proteins per sample, compared to the broader coverage achieved in our study. This comparison highlights the potential benefits of the enhanced sensitivity and resolution provided by integrating Mag-Net enrichment with Orbitrap Astral mass spectrometry.

### Validation of biomarkers confirms robustness and predictive value via AI modeling

To translate our deep proteomic findings into actionable clinical tools, we focused on the strong association between sampling interval and PC1 observed in our multivariate analysis (**Fig. 2B-C**). We prioritized the proteins driving this PC1-associated variance (35%), selecting candidates based on a stringent intersection of high FC rank and established biological relevance to PD pathophysiology. This selection process yielded a targeted list of 9 proteins, including DRG1, SCAP, and EMC4 (**Fig. 2F**). Additionally, we strategically included APLP1 and FN1 as hypothesis-driven candidates: APLP1 was selected for its mechanistic link to alpha-synuclein transmission^72^, while FN1 was included to clarify conflicting reports regarding its prognostic utility and potential role in fibrosis-associated neurodegeneration in PD^73^.

To establish a robust PD biomarker prediction model, we implemented a comprehensive machine learning benchmarking framework (**Fig. 3A**). We trained 3 logistic regression algorithms on all possible combinatorial panels consisting of four to six proteins selected from the nine candidates identified in the discovery cohort. Among the diverse algorithms and panels tested, over 60% (202 out of 336 combinations) of protein combination panels demonstrated superior performance, achieving an Area Under the Receiver Operating Characteristic (ROC) Curve (AUC) value of >0.9 (**Fig. 3B**). Furthermore, over 95% (319 out of 336 combinations) of panels demonstrated acceptable performance, achieving an AUC value of >0.7. This probabilistic approach is particularly well-suited for capturing non-linear biological relationships in clinical datasets. Notably, DRG1 and SCAP were most frequently selected in the top-performing panels, suggesting they serve as “anchor” features with high predictive stability for PD progression. These models significantly outperformed random guessing (AUC=0.5), confirming that our selected plasma signatures harbor genuine predictive value for distinguishing disease progression states from baseline diagnosis.

**Figure 3.**
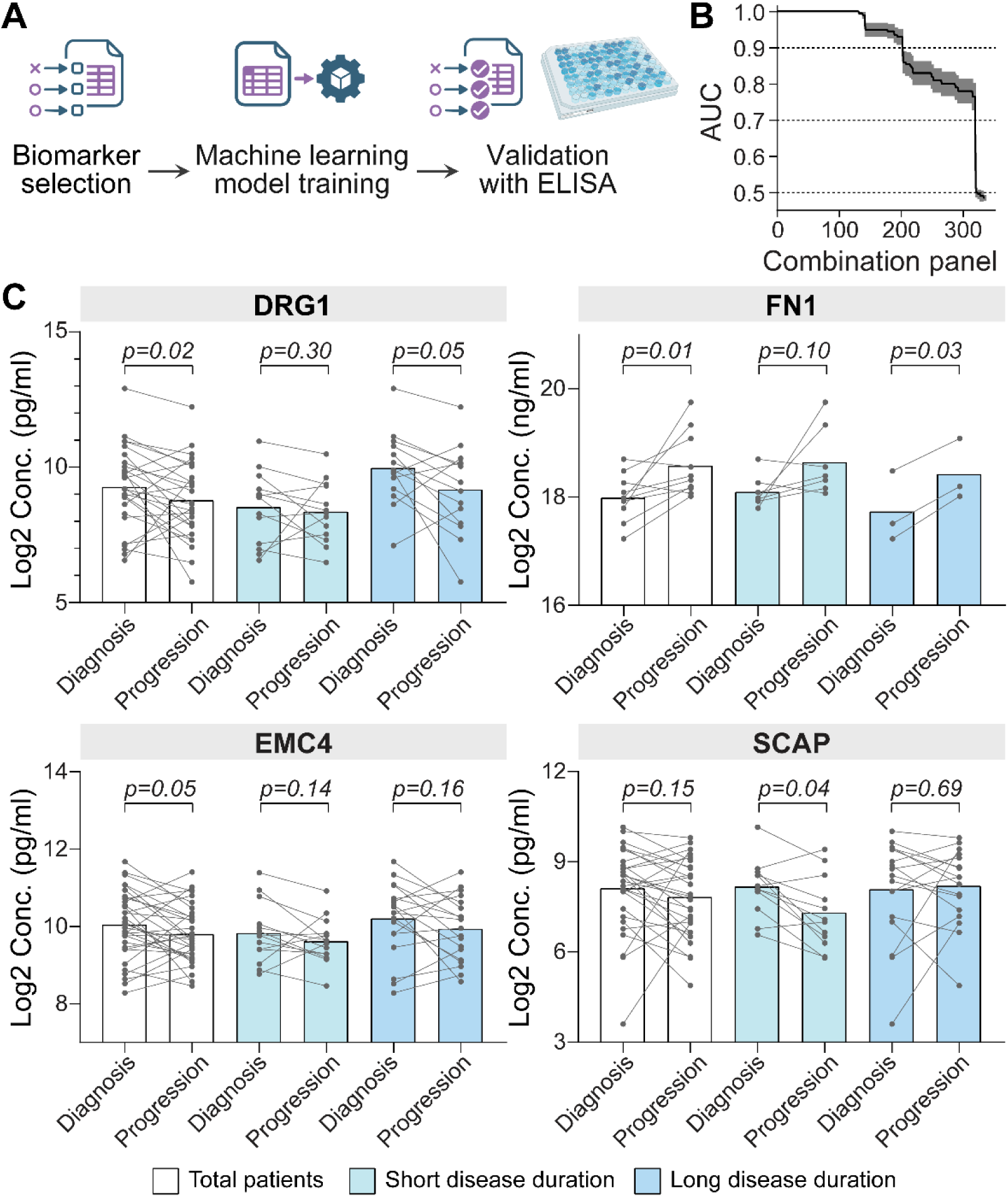
AI-driven prediction modeling and independent validation of candidate plasma biomarkers. **A.** Schematic flow for biomarker validation with machine learning-based prediction, created with BioRender. **B.** Performance benchmark of all protein combinations with logistic regression algorithms based on the evaluation metric ROC AUC. **C.** Sandwich ELISA showing plasma concentration of indicated proteins in diagnosis and progression samples obtained from Validation cohorts. Comparison between diagnosis and progression group (paired dataset) was analyzed using the paired t-test (two-tailed, empirical permutation). Illustration of paired data by connecting the data points.

To rigorously validate these findings in an independent population, we collected longitudinal plasma samples from a separate validation cohort of 38 PD patients (**Supplementary Table 3**). The validation cohorts were ensured to have a comparable distribution of key clinical demographics to those of the discovery cohort (**Supplementary Table 4**). We quantified plasma protein levels using an orthogonal method of sandwich ELISA (**Fig. 3C and Supplementary Fig. S3**). Consistent with our discovery proteomics data, FN1 and DRG1 exhibited significant differential abundance between diagnosis and progression timepoints in the total cohort (p=0.01 and p=0.02, respectively), while EMC4 showed a strong trend toward significance (p=0.05). Intriguingly, SCAP displayed a dynamic, time-dependent temporal pattern: while its levels remained stable when analyzing the cohort as a whole, it was significantly altered (p=0.05) specifically in the subset of patients with short sampling intervals. While the results of APLP1, VWA8, and GPAT4 were consistent with proteomics data, the difference is not significant; those of SRSF3 and APAF1 were inconsistent with the proteomics data. This independent corroboration statistically validates the multivariate signatures identified in our discovery phase (**Fig. 2**) and biologically substantiates the temporal dynamics of our candidates. Specifically, it reinforces the conclusion that critical pathogenic drivers, such as SCAP-mediated lipid metabolism, are not statically dysregulated, but rather transiently modulated during specific, biologically active windows of disease progression.

### Functional genomics for alpha-synuclein PFF uptake identifies novel biomarker candidates and potential drug targets

Our multivariate analysis established disease progression as a dominant factor shaping the plasma proteome (35% of variance explained by PC1 in **Fig. 2B&C**). To dissect the molecular drivers of this trajectory, we integrated plasma protein profiling with a functional genomics approach targeting alpha-synuclein PFF uptake. We reasoned that because PD progression is biologically defined by the spatiotemporal spread of pathological alpha-synuclein seeds across brain regions, their cellular uptake constitutes a critical rate-limiting step in disease initiation and progression. The internalization of PFFs by dopaminergic neurons acts as a critical priming event for PD pathogenesis^74,75^, initiating a prion-like cascade that induces misfolding and aggregation of endogenous α-synuclein, ultimately leading to cytotoxicity^76,77^. Crucially, this intracellular toxicity drives the secretion and shedding of stress-response proteins, like APAF1, a regulator of PFF-driven apoptosis^78^ that is upregulated in the plasma of PD patients, and CSF1R^78^, a receptor mediating neuroinflammation that is similarly elevated in peripheral blood. Additionally, LRRK2, a well-studied PD-associated kinase, modulates Rab-dependent endocytosis and endolysosomal trafficking, and pathogenic activation of this pathway impairs alpha-synuclein clearance^79^, thereby exacerbating PFF-driven seeding and toxicity^80^. Furthermore, cell-surface proteins such as APLP1^81^ have been directly implicated in the transmission of pathologic alpha-synuclein, linking cellular uptake mechanisms to detectable biofluid signatures. Thus, we hypothesized that the plasma proteome remodeling observed over time is, in part, a peripheral echo of this uptake-dependent propagation, making the molecular machinery governing PFF uptake a strategic target for intercepting disease initiation and progression.

To systematically dissect the genetic drivers of alpha-synuclein PFF uptake, we conducted a proteome-wide functional genomics screen using the **BOGO** (**B**xb1-landing pad human **O**RFeome-integrated system for a proteome-wide **G**ene **O**verexpression)^28^ platform based on the HeLa cell line (HeLa-ORFeome; **Fig. 4A**). Although the pathological sequelae of alpha-synuclein aggregation are neuron-specific, the initial cellular entry of fibrils relies on evolutionarily conserved endocytic mechanisms, such as macropinocytosis and heparan sulfate proteoglycan interactions, that operate across diverse cell types^29,82,83^. Leveraging this conservation, we utilized HeLa cells as a robust, high-throughput system to profile uptake regulators independent of cell-type-specific confounds. Following 16 hours of incubation with pH-sensitive PFFs (PFF-pHrodo)^84,85^, cells were sorted into populations exhibiting the lowest (bottom 10%) and highest (top 10%) uptake rates. ORFeome-sequencing^28^ revealed distinct overexpression profiles for each population (**Supplementary Fig. S4A and Supplementary Table S5**), with pathway analysis highlighting ‘Dopaminergic synapse,’ ‘Cholinergic synapse,’ ‘PI3K-Akt,’ and ‘AMPK’ signaling pathways, reinforcing their established roles in PD etiology^86–88^ (**Supplementary Fig. 4B**). To validate novel hits, we selected candidates not previously linked to PD, including DERL1, FOXO3, IFNG, LAMC1, and SULF2. Consistent with the primary screen, flow cytometry confirmed that overexpression of DERL1, FOXO3, IFNG, and LAMC1 significantly suppressed PFF uptake, whereas SULF2 overexpression significantly enhanced the uptake (**Fig. 4C**).

**Figure 4.**
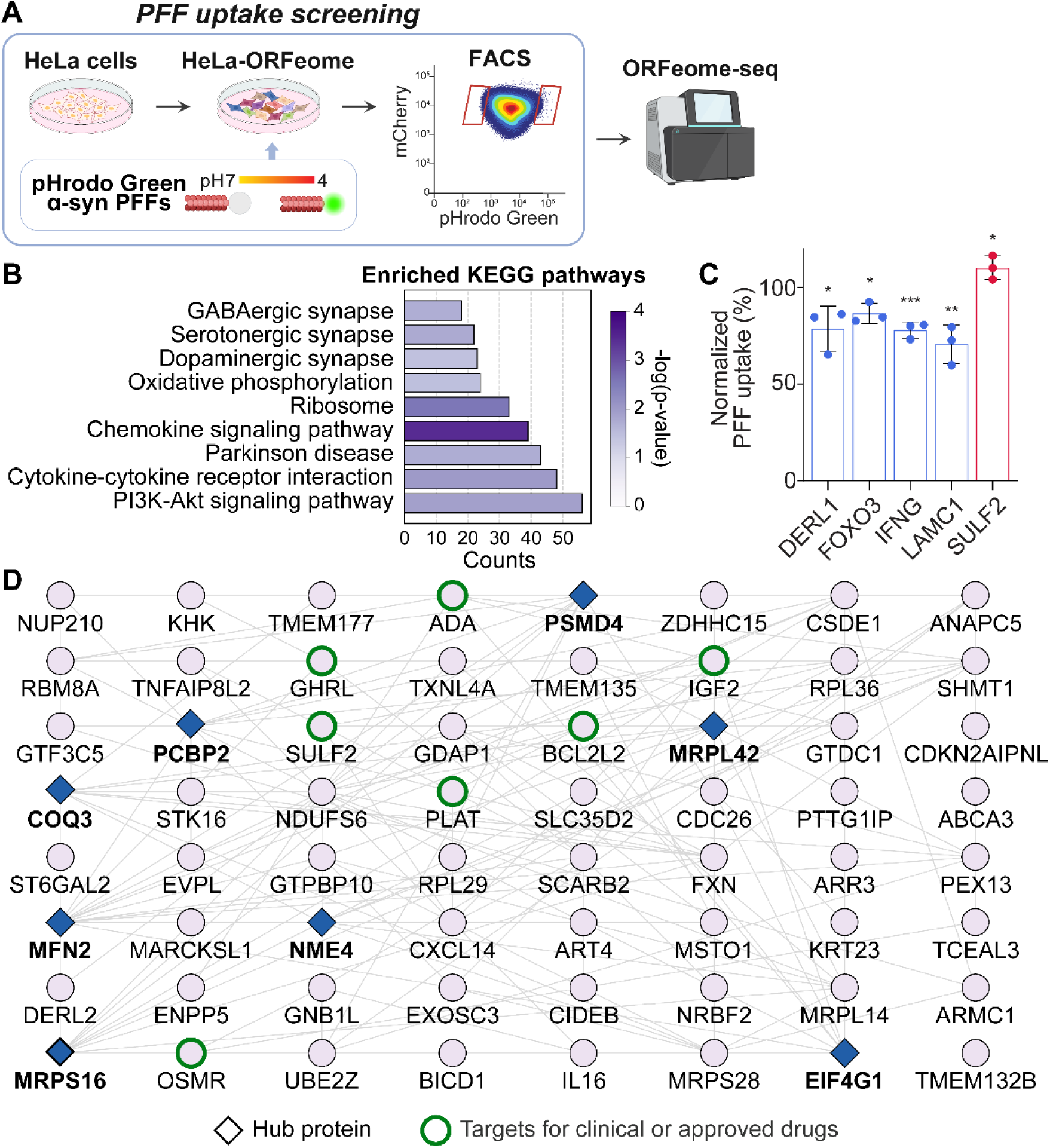
Identification of PFF uptake modulators and cross-validation with plasma signatures. **A.** Schematic illustration of PFF uptake screening, created with BioRender. **B**. KEGG pathway enrichment analysis of differentially represented ORFs from PFF uptake screening. **C.** Bar plot showing the relative pHrodo-positive population of hits identified by PFF uptake screening. P-value was calculated by two-tailed t-test. **D.** Subnetwork of 64 proteins showing significant changes in both the plasma proteomics data and PFF uptake screening data. Diamond-shaped nodes indicate hub proteins. Green-highlighted nodes denote proteins with clinical or approved therapeutic targets.

To identify plasma biomarkers involved in this mechanism, we intersected the PFF uptake screen hits with the top 1,000 DEPs discriminative between sampling intervals. This overlap yielded 64 candidate proteins (**Supplementary Fig. S4C and Supplementary Table S6**), which were mapped into a functional network (**Fig. 4D**). Five of the eight major hubs, PSMD4, MRPS16, MFN2, NME4, and EIF4G1, emerged and connected systemic plasma changes to core PD mechanisms. PSMD4, a proteasome subunit, links proteostasis failure to alpha-synuclein accumulation^89^, and MRPS16 is a crucial component of the small subunit of the mitochondrial ribosome linked to mitochondrial protein synthesis^90^. MFN2 mediates mitochondrial fusion and PINK1/Parkin mitophagy^91^, while NDUFS6 reflects oxidative phosphorylation deficits^92^. NME4 is a mitochondrial protein linked to clearing damaged mitochondria^93^, and EIF4G1 is implicated in familial PD via translational control^94^. We also identified SCARB2^95^, GDAP1^96^, ZDHHC15^97^, and PLAT^98^ as functionally relevant candidates. Importantly, PSMD4^99^, and PLAT^100^ are targets of approved drugs, highlighting translational opportunities for repurposing existing therapies (**Fig. 4D**).

### Functional genomics for alpha-synuclein PFF-induced toxicity with dopamine-positive neurons provides more physiologically relevant biomarker candidates with therapeutic potency

To further validate these findings in a model with greater physiological relevance to PD, we investigated driver genes associated with neuroinflammation and toxicity in dopamine-positive neurons. We applied the **LOGO** (**L**entivirus and **O**RFeome integrated proteome-wide **G**ene **O**verexpression perturbation)^28^ system in human H9 pluripotent stem cells (hESC; **Fig. 5A**). Following differentiation into dopamine-positive neurons and subsequent doxycycline treatment for gene expression activation, cells were challenged with PFFs and analyzed via ORFeome-sequencing^28^. The survival profiles of the pre- and post-treatment groups were clearly segregated, underscoring the robustness of the assay (**Supplementary Fig. 4B**). Notably, this analysis independently re-identified the ‘PI3K-Akt,’ and ‘AMPK’ signaling pathway as a critical determinant of neuronal response (**Fig. 5B**), corroborating the mechanisms observed in our primary screen. Moreover, genes related to cellular senescence, which have been linked to dopaminergic neuron loss in PD mouse models^101^, were also enriched.

**Figure 5.**
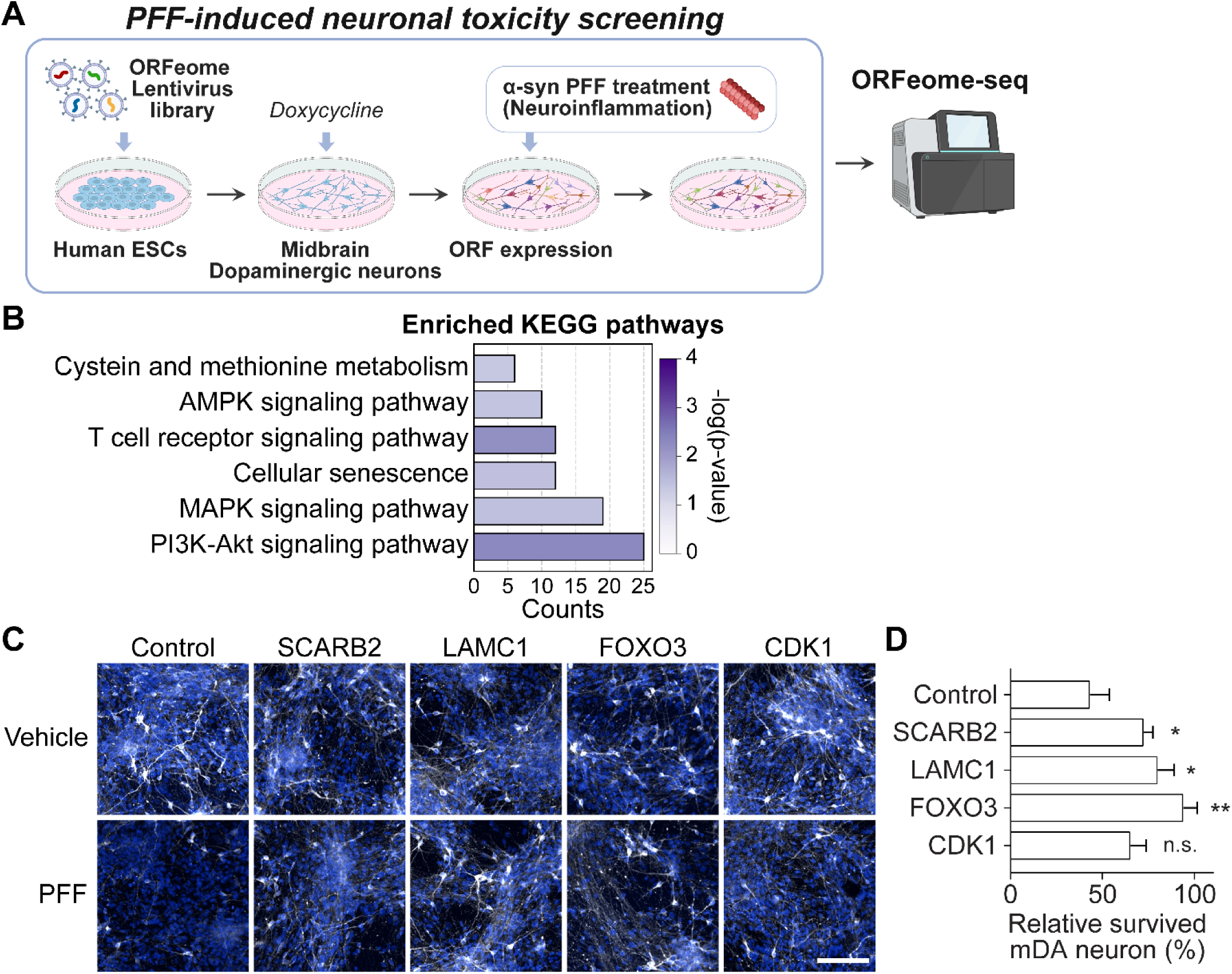
Identification of PFF-induced neural toxicity modulators through functional genomics screening in dopaminergic neurons. **A.** Schematic illustration of PFF-induced neuronal toxicity screening, created with BioRender. **B**. KEGG pathway enrichment analysis of differentially represented ORFs from PFF-induced toxicity screening. **C.** Survived dopaminergic neurons were assessed by confocal microscopy, measuring tyrosine hydroxylase (TH, white) and Hoechst-stained nuclei (blue) in Vehicle or PFF treatment with 5 μg/ml of PFF for 14 days. Scale bar, 100 μm. **D.** Bar plot showing the relative survival of dopaminergic neurons with overexpression of hits identified by PFF-induced toxicity screening. P-value was calculated by a one-tailed t-test.

To further validate hits that were also significant in the PFF uptake screen (**Fig. 4**), we selected 4 candidates, SCARB2, LAMC1, FOXO3, and CDK1. Consistent with the primary screen, overexpression of SCARB2, LAMC1, and FOXO3 significantly rescued dopaminergic neurons from the PFF-induced toxicity, whereas CDK1 overexpression had a negligible effect on neuronal survival following PFF treatment (**Fig. 5C and D**). We observed a statistically significant overlap (p<0.001, Fisher’s exact test) between two PFF-related functional genomics screens (**Supplementary Fig. 4C**). Especially, both FOXO3 and OSMR were enriched in neurons that survived PFF exposure (**Supplementary Table S5 and S6**). Interestingly, each factor is known to exert context-dependent, dual roles in the nervous system^102–104^. These screening results suggest that upregulation of FOXO3 and OSMR in dopaminergic neurons may mitigate PFF-induced neuroinflammation and PD progression.

Finally, we integrated these functional insights with clinical genetics to bridge the gap between observational genomics and mechanistic validation^105^. We cross-referenced our data with a recent study identifying 137 PD-associated copy number variations (CNVs)^106^. We observed a statistically significant overlap (p<0.001, Fisher’s exact test) between these genetic hits and our proteomic profiles, with four proteins (PARK7, SNCA, RAB32, and VPS35) differentially expressed in patient plasma. Moreover, three of these markers (SNCA, LRRK2, and VPS35) were independently identified as regulators of PFF uptake in our functional screen with significant overlap (p<0.001). Additionally, we analyzed two published snRNA-seq datasets^107,108^ of midbrain tissues from healthy controls and PD patients. In dopaminergic neurons, 105 differentially expressed genes (DEGs) were shared between the two datasets (|log_2_FC|>0.58 and p<0.05). Among them, 10 genes were identified as DEPs in plasma proteomics data as well. Especially, PLOD3^109^ and KIT^110^ were both upregulated in Progression samples of plasma proteomics data and in Patient samples of snRNA-seq data, as previous studies have implicated these genes in PD and neurodevelopmental processes, respectively. This convergence of plasma proteomics, functional screening, and human genetics underscores the biological accuracy of our platform.

### Finding novel biomarkers for response against dopaminergic treatment through meta-analysis

Our multivariate analysis identified LEDD as one of the primary determinants of plasma proteomic remodeling (12% of variance explained by PC2 in **Fig. 2B, D**). However, because plasma inherently captures systemic signals from peripheral tissues, distinguishing biomarkers of central therapeutic response from peripheral metabolic effects remains a challenge. To deconvolute this complexity and isolate biomarkers with direct relevance to CNS responsiveness, we employed a cross-species, cross-omics validation strategy.

We reasoned that intersecting our plasma protein signatures with brain-specific transcriptomes would act as a ‘mechanistic filter,’ prioritizing candidates that trace back to gene expression changes within dopaminergic circuits. To this end, we compared our LEDD-dependent targets with transcriptomic datasets from three *in vivo* studies with mouse models, profiling L-DOPA effects directly in the neural microenvironment (**Supplementary Fig. 5**). These included: i) cell-type-specific L-DOPA responses in striatonigral and striatopallidal medium spiny neurons^111^ (GSE291024); ii) bulk transcriptional changes in the mouse brain following L-DOPA treatment (unpublished; GSE279704); and iii) spatial transcriptomic profiling of the forebrain in a model of progressive dopaminergic loss^112^ (Zenodo: 14762864).

By integrating our putative LEDD biomarkers with DEGs in the mouse brain, we identified 42 candidates potentially involved in L-DOPA responsiveness in PD. Among these, four genes of NDUFS4, GNAS, TSC1, and NTS emerged as particularly relevant to dopaminergic treatment response. NDUFS4, a core component of mitochondrial complex I, represents a critical link between cellular bioenergetics and dopamine metabolism, consistent with the established role of mitochondrial dysfunction^113^ in PD pathophysiology. GNAS, encoding a G-protein alpha subunit, directly influences dopamine receptor signaling through cAMP-mediated pathways^114^, suggesting that genetic variation may modulate drug sensitivity. Similarly, TSC1, a regulator of mTOR signaling^115^, is noteworthy given the interplay between mTOR activity and dopamine receptor function^116^. Finally, NTS acts as an intrinsic modulator of dopaminergic circuits by regulating dopamine release and receptor sensitivity^117^. In addition, four other candidates of IDH2^118^, DNASE1^119^, ABCA3^120^, and HNRNPH2^121^ are already targeted by approved or investigational therapies. These examples underscore the translational potential of our findings and highlight opportunities for leveraging targeted approaches to complement dopaminergic therapies in PD.

### Integrative network analysis of our novel clinical biomarkers and functional candidates

To synthesize insights from our multi-layered approach, we constructed an integrative network (**Fig. 6A**) combining (i) DEPs associated with disease progression obtained from plasma proteomics (**Figs. 2 and 3**), functional candidates identified through (ii) PFF uptake screens (**Fig. 4**), and (iii) PFF-induced toxicity screens (**Fig. 5**). This network was generated using high-confidence protein–protein interaction data^122^ and visualized to highlight functional modules and hub nodes.

**Figure 6.**
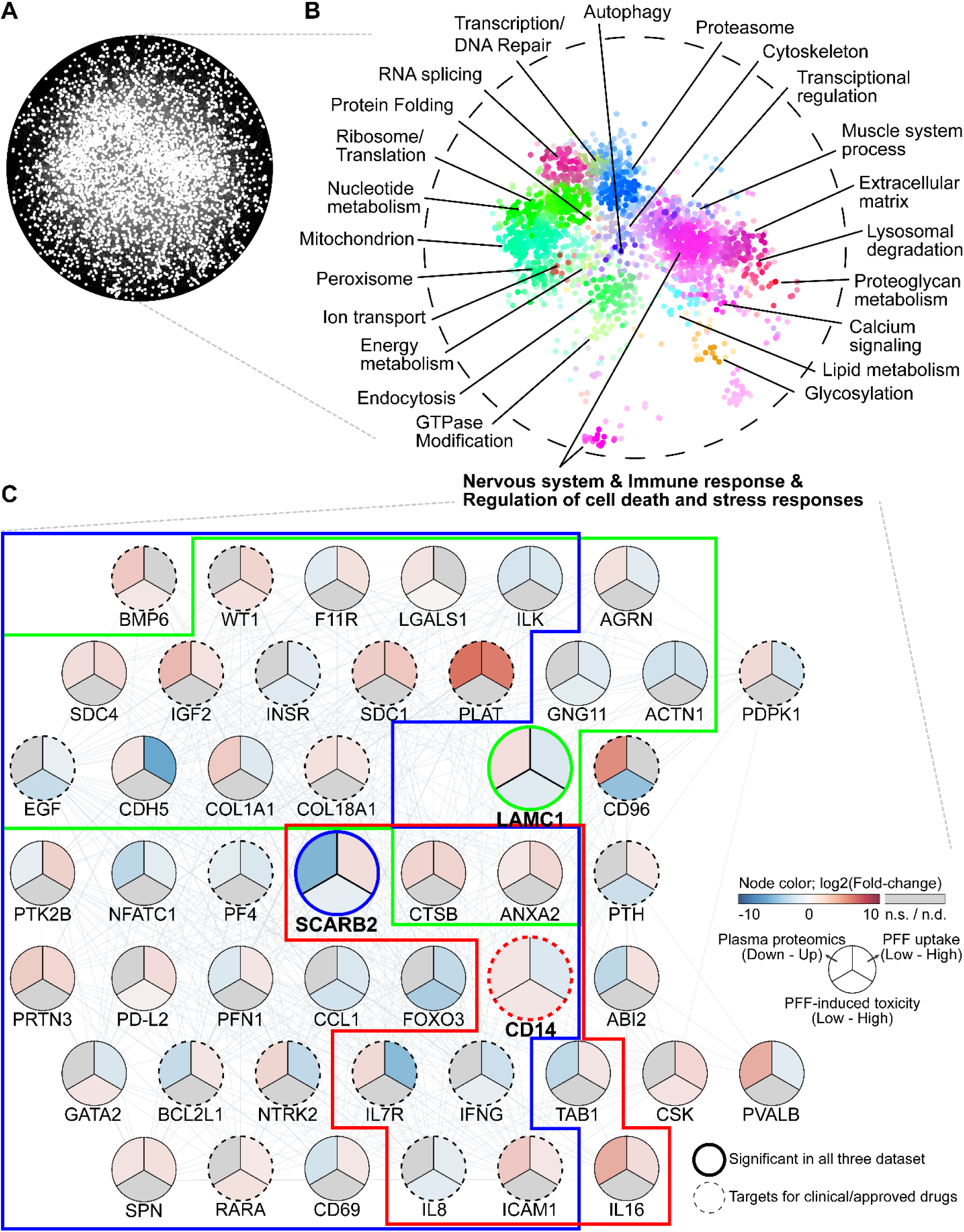
Integrative network analysis of clinical plasma biomarkers and functional genomic candidates. **A.** Protein-protein interaction network integrating disease progression-associated DEPs from plasma proteomics with candidates identified from PFF uptake and PFF-induced toxicity screens. **B.** SAFE-based functional enrichment map highlighting significantly overrepresented pathways. **C.** Subnetwork of 48 proteins showing significant changes in at least two datasets and high neighborhood scores. Node colors indicate log_2_ fold changes across plasma proteomics, PFF uptake, and PFF-induced toxicity datasets. Boxed regions highlight subnetworks centered on the hub proteins SCARB2, LAMC1, and CD14. Dotted nodes denote proteins with clinical or approved therapeutic targets.

Functional enrichment of the network confirmed significant overrepresentation of pathways implicated in PD, including muscle system process, mitochondrion, and proteostasis (**Fig. 6B**). These pathways showed substantial overlap with those identified in our PFF–related screening analyses. Among the identified clusters, one module showed strong enrichment for pathways related to nervous system development (**Pink central cluster in Fig. 6B**). Subsequent analysis focused on a subnetwork of 48 proteins exhibiting significant changes in at least two datasets with a maximum neighborhood score, within which LAMC1, CD14, and SCARB2 emerged as consistently altered across all three datasets (**Fig. 6C**). Of particular interest are CD14^123^ and SCARB2^124^; although previously identified as genetic risk factors for PD in population-wide studies, the molecular mechanisms underlying their contribution to disease have been unclear. Our findings resolve this ambiguity, highlighting the capacity of this integrative framework to decode the functional biology behind clinical genetic associations. Furthermore, CTSB and ANXA2 were connected to these three proteins through experimental interaction evidence or co-expression relationships. Both proteins are functionally linked to proteostasis and vesicular transport processes that have been increasingly associated with neurodegenerative pathology^125,126^.

Finally, to explore the translational potential of these findings, we overlaid Drug-Gene Interaction database information^127^ onto our network (**Fig. 6C, dotted nodes**). This analysis identified several “druggable” nodes among our validated candidates. Notably, we found that approved pharmacological agents exist for targets such as PLAT^100^, NTRK2^128^, and PTH^129^. The identification of these druggable targets within key pathological hubs provides a rationale for repurposing existing therapies to potentially modify disease progression or complement current dopaminergic treatments.

Collectively, this integrative network provides a systems-level view of how clinical biomarkers and functional candidates converge on core biological processes driving PD progression. These findings offer a prioritized set of targets for mechanistic validation and translational development.

## Discussion

The absence of accessible, robust biomarkers for PD progression has long impeded the development of disease-modifying therapies^130^. While CSF offers a closer approximation of CNS pathology, its invasive nature restricts its utility for routine longitudinal monitoring^131^. Plasma represents an ideal alternative, yet historical efforts have been stymied by the “dynamic range problem,” where high-abundance proteins mask the low-abundance, brain-derived signals essential for monitoring neurodegeneration^132^. In this study, we overcame this barrier by integrating Mag-Net^25^-based EV enrichment with Orbitrap Astral mass spectrometry^26,27^. This approach achieved an unprecedented depth of ∼6,500 plasma proteins, a coverage magnitude higher than conventional datasets^39–41^, allowing us to deconvolute the complex systemic signatures of PD into two actionable axes: intrinsic disease progression and pharmacological response.

A major finding of this work is the rigorous separation of proteomic changes driven by disease duration from those induced by dopaminergic therapy. Previous cross-sectional studies^39–41^ comparing patients to healthy controls have often struggled to disentangle these factors. By employing a longitudinal design with paired samples, coupled with multivariate modeling, we identified that LEDD is one of the primary drivers of plasma proteomic variance, distinct from the disease trajectory itself. Our meta-analysis of L-DOPA-treated animal models validated this distinction, pinpointing proteins such as NDUFS4, GNAS, TSC1, and NTS as specific markers of dopaminergic response. This segregation is critical for clinical trial design, as biomarkers intended to track neuroprotection must be insulated from the proteomic noise generated by symptomatic relief medications. Conversely, the LEDD-responsive panel offers a potential readout for titrating therapy and monitoring patient compliance or resistance.

Beyond biomarker identification, our study bridges the gap between systemic observations and central disease mechanisms through functional genomics. A recurring criticism of plasma proteomics is the uncertainty regarding the tissue of origin and the pathological relevance of identified proteins^133^. We addressed this by intersecting our progression-associated plasma signatures with a proteome-wide ORFeome screen for regulators of PFF uptake and cytotoxicity. This convergence highlighted a specific network of proteins, including LAMC1, CD14, and SCARB2, that are not only detectable in plasma but functionally drive the cell-to-cell transmission of alpha-synuclein pathology.

The translational potential of these findings is underscored by our identification of “druggable” hubs within these pathogenic networks. Our network analysis revealed that central drivers of progression identified here, such as NTRK2 and PLAT, are targets of approved pharmacological agents. This suggests that the plasma biomarkers identified in our deep-profiling workflow could serve dual roles: as surrogate endpoints for clinical trials and as direct targets for drug repurposing strategies aimed at stabilizing lysosomal function or mitochondrial dynamics. Furthermore, the successful validation of select candidates via ELISA and their incorporation into AI-driven predictive models demonstrates the feasibility of translating these high-dimensional discoveries into accessible clinical assays.

Our study has limitations that warrant consideration. While our longitudinal design minimizes inter-individual variability, the discovery cohort size (n=10) is relatively small. Although we validated key targets in an independent cohort (n=38) and leveraged large-scale public datasets for cross-validation, replication in multi-center cohorts with diverse genetic backgrounds is necessary to establish universal applicability. Additionally, while Mag-Net enriches for EV-associated proteins, plasma remains a peripheral biofluid; future multi-omics studies integrating paired CSF analysis to bridge peripheral and central signatures, PET imaging, or post-mortem tissue analysis would further refine the correlation between these peripheral signals and specific regional neurodegeneration.

In summary, this study provides a systems-level resource that fundamentally expands the observable plasma proteome in PD. By synergizing ultra-deep proteomics with proteome-wide functional genomic screening, we have moved beyond simple cataloging to identify mechanistic drivers of disease progression and treatment response. These findings lay the groundwork for a new generation of precision medicine tools capable of stratifying patients, monitoring therapeutic efficacy, and guiding the development of disease-modifying interventions.

## Acknowledgment

This research was supported by a grant from the Korea Health Technology R&D Project through the Korea Health Industry Development Institute (KHIDI), funded by the Ministry of Health & Welfare, Republic of Korea (RS-2023-00265820); the Bio & Medical Technology Development Program through the National Research Foundation (NRF), funded by the Ministry of Science & ICT, Republic of Korea (RS-2024-00411768, RS-2024-00440778, RS-2024-00332454); Global -Learning & Academic Research Institution for Master’s·PhD students, and Postdocs (LAMP) Program of the NRF grant funded by the Ministry of Education (RS-2024-00445180); and a startup grant from Montreal General Hospital Foundation and Aune Foundation, and a Tomlinson award from McGill University. The Biospecimens and data used for this study were provided by the Biobank of Seoul National University Hospital, a member of the Korea Biobank Network (project No. 2024ER050800).

## Author Contribution

**J.-S.K., C.J., D.-K.K.,** and **H.-J.K.** conceptualized and designed the study. **J.-S.K.** performed the experiments, processed the data, analyzed the results, drafted the original manuscript, managed the project, and prepared the figures. **C.J.** performed the experiments and drafted the original manuscript. **M.S.K.** performed PFF-induced toxicity screening. **H.K.** carried out data curation, figure editing, and critical manuscript review. **S.L., S.H.H.,** and **K.A.W.** prepared the samples from two independent patient cohorts. **W.L.,** and **I.S.** provided critical resources. **S.N.K., B.L.T., S.K.,** and **C.A.B.** performed plasma proteomics. **K.B.J., S.S., E.L.**, and **Y.L.** assisted with data processing and manuscript editing. **A.G.C.**, **M.L.**, **G.L.**, **J.S.**, **G.P.**, **N.S.**, **J.-H.C.**, **J.-H.P.**, **D.E.H.**, **H.L.**, **K.A.M.**, **J-F.T.**, **J.-H.S.**, **J.R.**, and **T.D.** contributed to conceptualization, provided resources, experimental advice, manuscript editing, and technical support. **K.-J.Y**., **D.-K.K.,** and **H.-J.K.** supervised the project, provided conceptual guidance, and oversaw project administration.

## Supplementary Figures

**Supplementary Figure S1.**
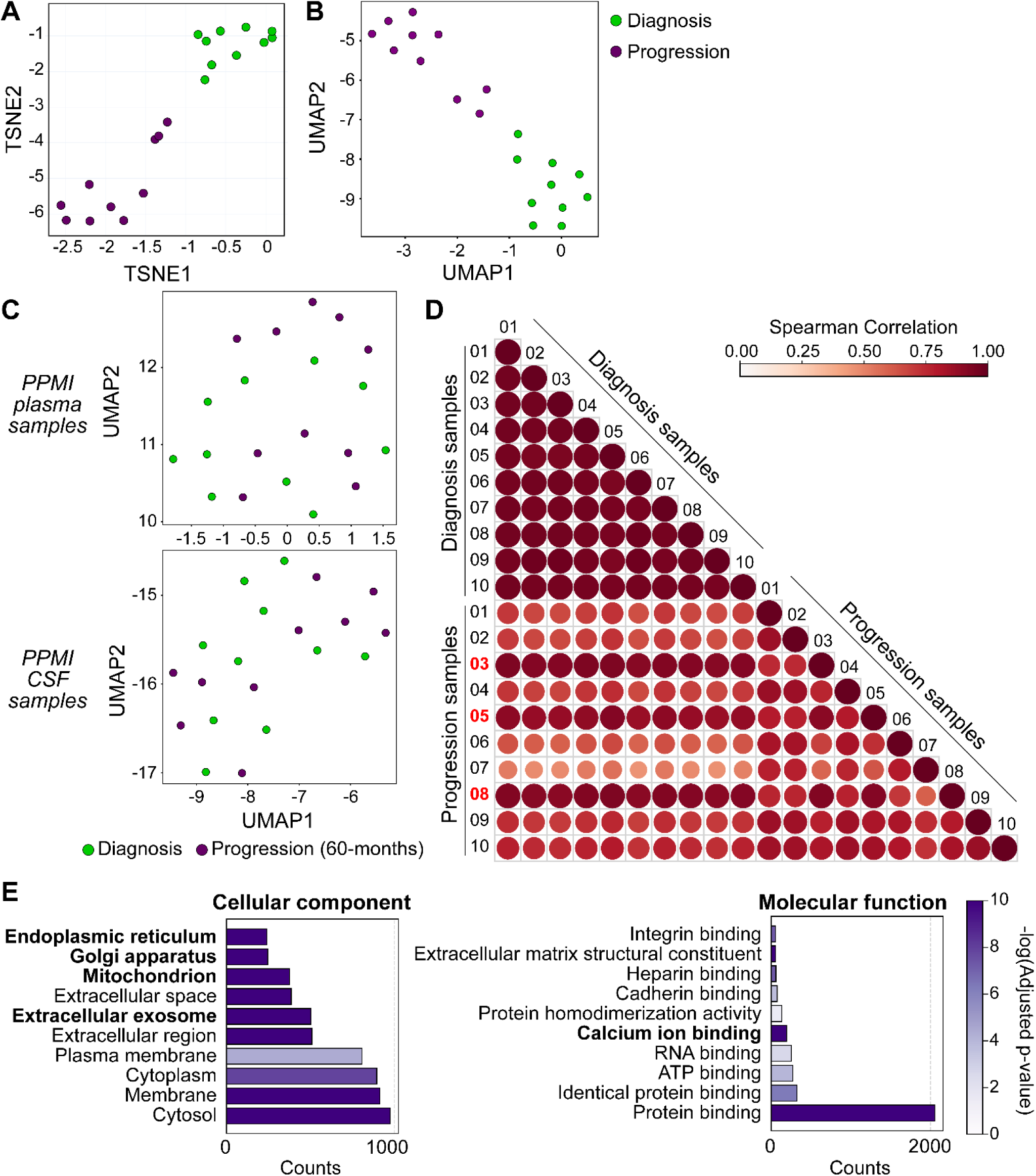
Overview of plasma proteomic profiling and comparison with external datasets. **A.** t-SNE plot of our longitudinal plasma proteomics dataset. **B.** UMAP analysis for comparison of the number of identified plasma proteins. **C.** UMAP analysis of longitudinal plasma and CSF proteomics data obtained from the PPMI database. Progression samples were defined as those collected more than 60 months after diagnosis. **D.** Correlation matrix of longitudinal plasma proteome profiles showing higher intra-group than inter-group correlations, supporting reproducibility. Red indicates patients with short disease duration. **E.** Functional enrichment analysis of DEPs. The top 10 GO molecular function and cellular component terms were identified.

**Supplementary Figure S2.**
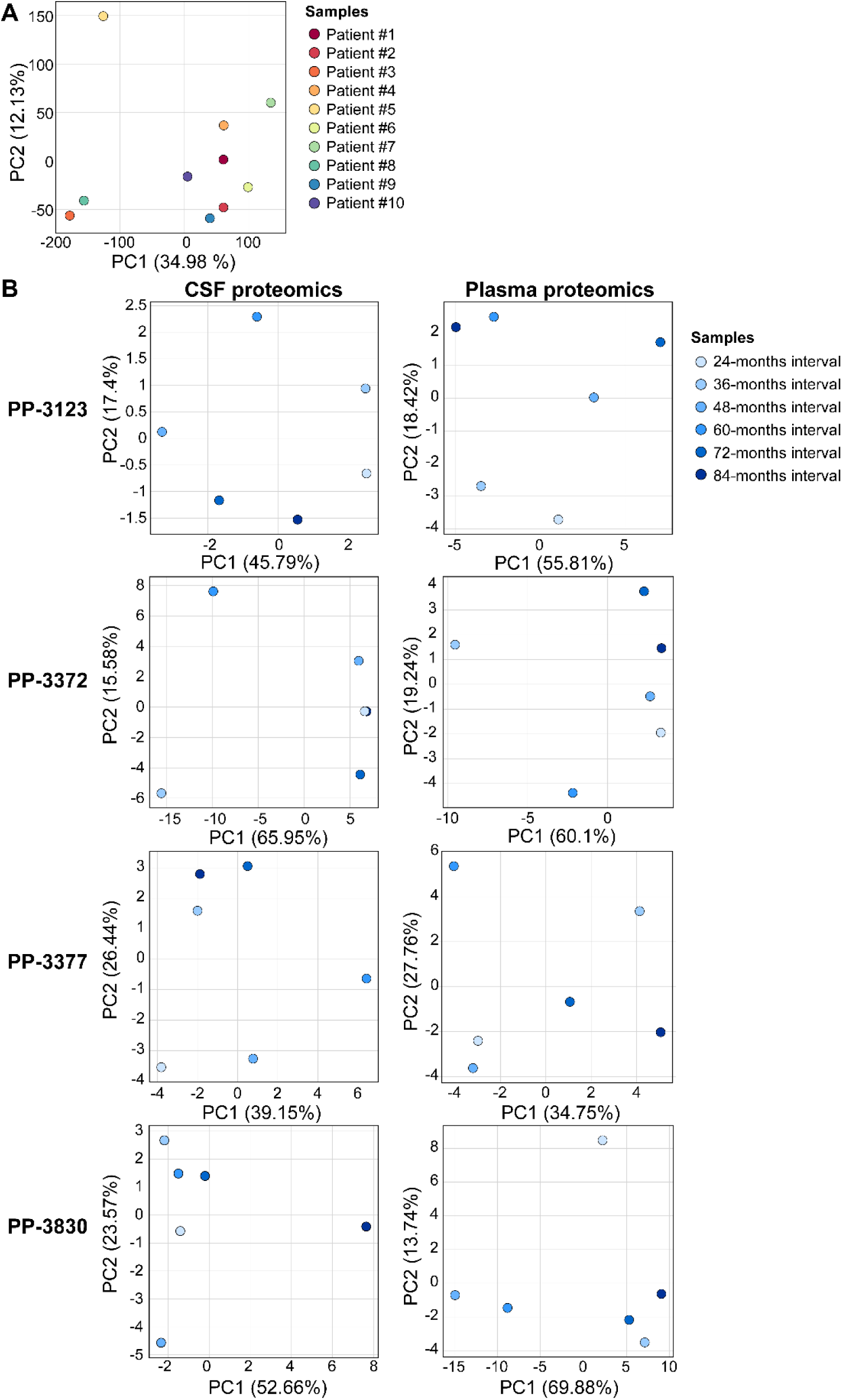
Principal component analysis of longitudinal proteomic changes. **A.** PCA plots of FCs for individual patients. **B & C.** PCA plots of FCs for progression sample versus diagnosis sample in several interval points from (**B**) CSF and (**C**) plasma samples of 4 different patients. The raw data of these samples were obtained from the PPMI database.

**Supplementary Figure S3.**
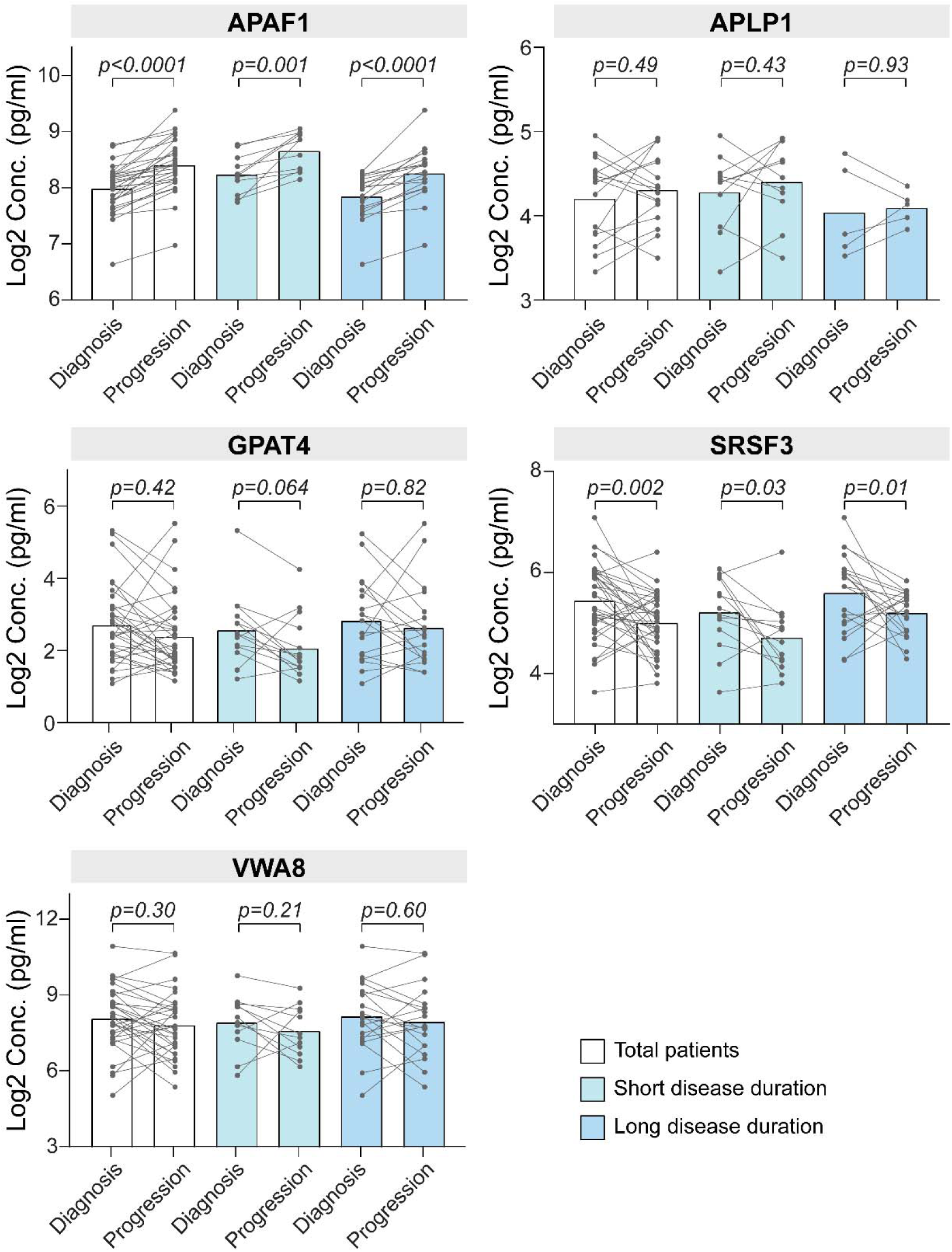
Validation of candidate biomarkers using ELISA. Sandwich ELISA quantification of indicated plasma proteins in paired diagnosis and progression samples from the independent validation cohort. Statistical significance was assessed using a two-tailed paired t-test with permutation. Lines connect data points from the same individual.

**Supplementary Figure S4.**
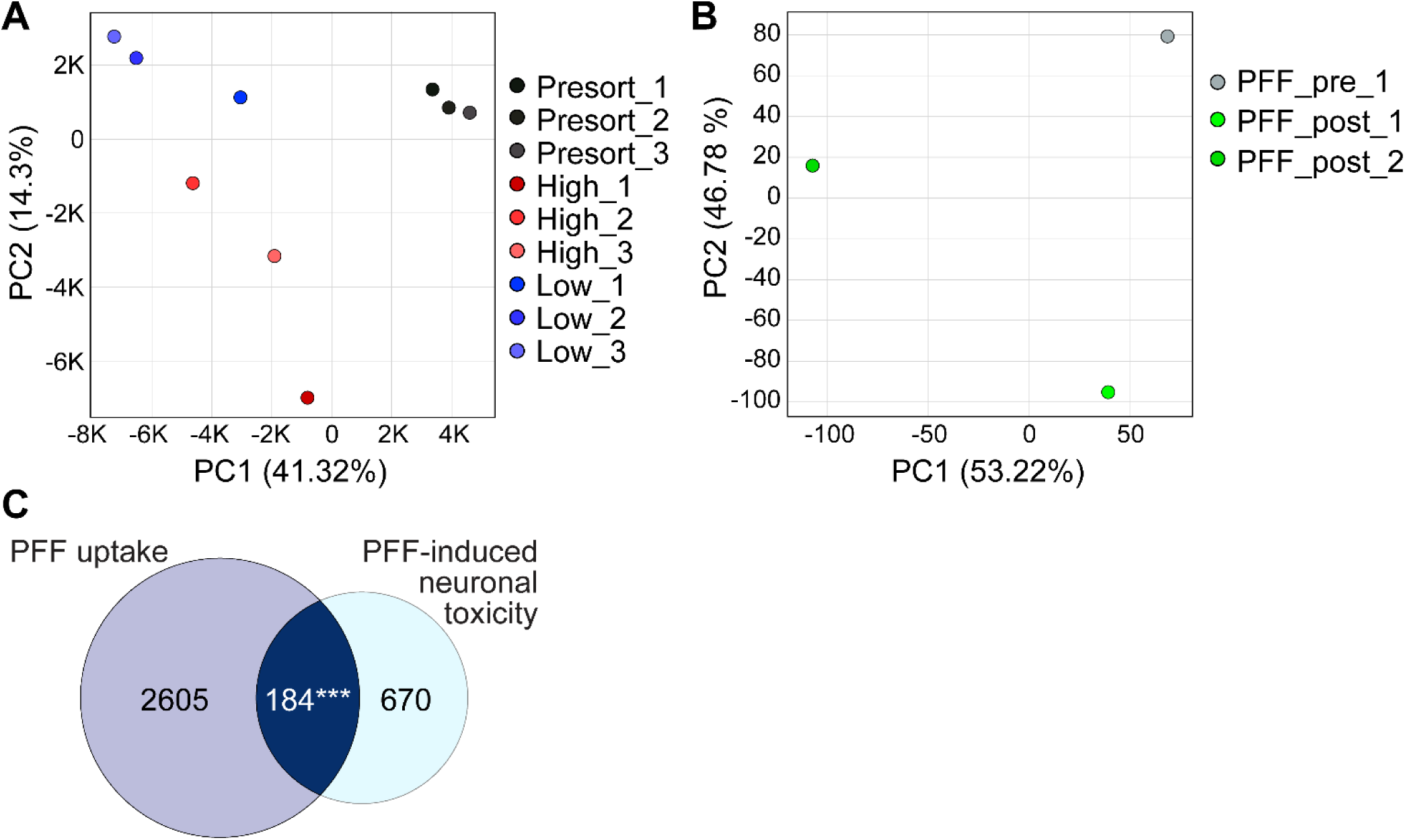
Quality control and functional enrichment of functional genomic screens. **A & B.** PCA plots of (**A**) PFF uptake screening and (B) PFF-induced toxicity screening results. **C**. Venn diagram showing the overlap of differentially represented ORFs between two PFF-related gene overexpression screening data. For PFF uptake screening, the differentially represented ORFs were identified |log_2_FC|>0.58 and p<0.05. For PFF-induced toxicity screening, the differentially represented ORFs were identified |log_2_FC|>0.32 and p<0.05.

**Supplementary Figure S5.**
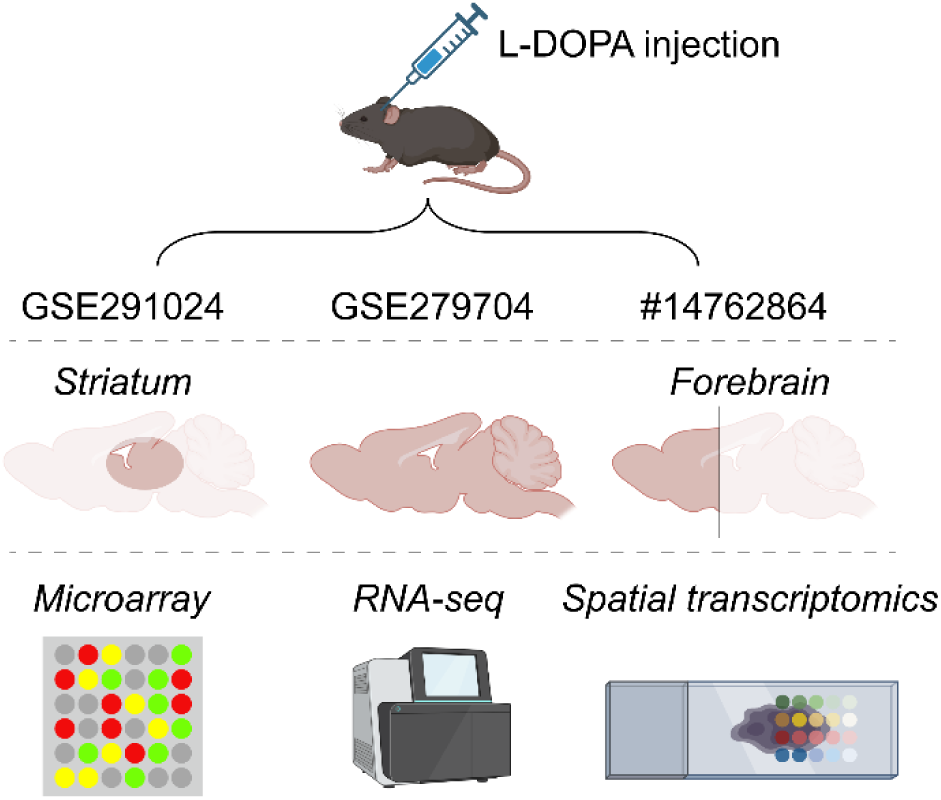
Schematic illustration of meta-analysis to identify genes that reacted against dopaminergic treatment.

## Supplementary Tables

**Supplementary Table 1.** Patient information of the Discovery cohort

**Supplementary Table 2.** Plasma proteomics result

**Supplementary Table 3.** Patient information of the Validation cohort

**Supplementary Table 4.** Clinical characteristics of the discovery and validation PD cohort in the study

**Supplementary Table 5.** PFF uptake screening

**Supplementary Table 6.** PFF-induced toxicity screening

## Methods and Materials

### Patient recruitment

We prospectively recruited ten drug-naïve Parkinson’s disease (PD) patients who were diagnosed between February 2017 and March 2018 at the outpatient clinic of the Department of Neurology, Seoul National University Hospital (IRB no. 2403-004-1515). Patients who had received prior pharmacological treatment for PD at baseline were excluded from the discovery cohort (**Supplementary Table 1**). After informed consent, baseline blood samples were collected in the drug-naïve state. Additional blood samples were obtained at a follow-up time point, ranging from 2 to 7 years after baseline.

### Patient information

Demographic data, including age and sex, along with clinical information such as the year of symptom onset, initial presenting symptoms, and medication status at follow-up, were collected. Sampling data, including the dates of baseline and follow-up sample collection, were also recorded. H-Y stage^134^ was assessed for disease severity. LEDD^135^ for each patient was calculated based on medication at follow-up sampling.

### Plasma collection from the patients

We dealt with patient blood as previously following the standard protocol of Seoul National University Hospital^136^. Whole blood samples (5 ml) were collected from each participant using ethylenediaminetetraacetic acid (EDTA) tubes. The tubes were gently inverted 10 times to ensure thorough mixing with the anticoagulant. Blood samples were then centrifuged at 1,500 × g for 15 min at RT. Following centrifugation, 4 ml of plasma was carefully aspirated using a pipette, ensuring minimal disturbance of the buffy coat and red blood cell layers. The plasma was then aliquoted into sterile containers and immediately stored at −80°C for long-term preservation.

### Membrane particle enrichment coupled with protein aggregation capture

Membrane particle capture from plasma and protein aggregation capture (PAC) steps were performed on a KingFisher Flex (Thermo Scientific). The combination of the membrane particle capture with PAC and digestion is referred to as Mag-Net^25^.

HALT cocktail inhibitor (Thermo Scientific, 78438) was added to plasma and then mixed 1:1 by volume with Binding Buffer (BB, 100 mM Bis-Tris Propane, pH 6.3, 150 mM NaCl). MagReSyn^®^ strong anion exchange or MagResyn^®^ Hydroxyl magnetic microparticles (ReSyn Biosciences) were first equilibrated 2 times in Equilibration/Wash Buffer (WB, 50 mM Bis-tris propane, pH 6.5, 150 mM NaCl) with gentle agitation and then combined in a 1:4 ratio (volume beads to volume starting plasma) with the plasma, followed by incubation for 45 min at RT. The beads were washed with WB 3 times for 5 min with gentle agitation. The enriched membrane particles on the beads were then solubilized and reduced in 50 mM Tris, pH 8.5, 1% SDS, 10 mM TCEP (Thermo Scientific, 20490) with 800 ng of standard enolase added as a process control. Following the reduction, the plate was removed from the Kingfisher. Samples were alkylated with 15 mM iodoacetamide (Sigma-Aldrich, A3221) in the dark for 30 min and then quenched with 10 mM DTT for 15 min. Unfractionated (“neat”) whole plasma (1 µl) was prepared in parallel (reduction, alkylation, and quenching) and added to the Kingfisher plate as a control for enrichment. The samples were processed using protein aggregation capture with minor modifications. Briefly, the samples were adjusted to 70% acetonitrile, mixed, and then incubated for 10 min at RT to precipitate proteins onto the bead surface. The beads were washed 3 times in 95% acetonitrile and 2 times in 70% ethanol for 2.5 min each on the magnet scaffold. Samples were digested for 1 hour at 47 in 100 mM ammonium bicarbonate with trypsin (Thermo Scientific #90057) at a ratio of 1:20 trypsin to protein. The digestion was quenched to 0.5% trifluoracetic acid and spiked with Pierce Retention Time Calibrant (PRTC) mixture (Thermo Scientific, 88321) to a final concentration of 50 nM. Peptide digests were then stored at −80 until LC-MS/MS analysis.

### Liquid chromatography and mass spectrometry

2 µl of each Mag-Net enriched sample with 50 nM of PRTC was loaded onto the Thermo Vanquish Neo HPLC coupled to the Thermo Orbitrap Astral mass spectrometer with a trap-and-elute workflow using an Acclaim PepMap 100, 75 μm x 2 cm for the trap column and an IonOpticks Aurora Elite C18, 1.7 μm, 75 μm x 15 cm column for the separation column. The separation column temperature was set to 55. Buffer A was 0.1% formic acid in water, and buffer B was 0.1% formic acid in 80% acetonitrile. For each sample, a 24-min separation gradient was used with a total injection-to-injection time of 30 min. The 24-min gradient starts at 4% B at 700 nl/min and has the following steps: 0.5 min at 4% B at 700 nl/min, 0.1 min to 8%B at 500 nl/min, 0.3 min at 8% B at 500 nl/min, 12.5 min to get from 8% to 22.5% B at 500 nl/min, 7.3 min to get from 22.5% to 35% B, and 0.4 min to get from 35% to 55% B at 700 nl/min. The column wash steps are 3 min at 99% B at 700 nl/min. Column equilibration was performed with Pressure Control at 800 bar with an equilibration factor of 4, and the Trap column used Fast Wash and Equilibration with Pressure Control at 800 bar and Zebra Wash using 4 Zebra Wash Cycles. Blanks were added between every 4 samples, with an extra injection added at the start of the samples to condition the column.

### DIA-MS using the Thermo Orbitrap Astral mass spectrometer

The system was operated at a full-MS resolution of 240,000 with a full scan range of 380–980 m/z. The full-MS AGC was set at 500%. Fragment ion scans were recorded at a resolution of 80,000 and maxIT of 3 ms. We used 300 windows of 2-Th scanning from 380 to 980 m/z with an MS2 scan range of 150-2000 m/z. The isolated ions were fragmented using HCD with 27% Normalized Collision Energy and collected in centroid mode.

### Analysis for protein quantification

Data analysis of all DIA files was performed using DIA-NN software (version 1.9.2)^137^. A search against the human proteome database from UniProtKB (downloaded in November 2024, 54,747 entries) was performed using a library-free workflow. Trypsin was used *in silico* digestion with a maximum allowance of 1 missed cleavage. The cutoff for the false discovery rates (FDRs) at the protein and peptide levels was set to 1%. A match between runs was allowed. Raw files were deposited in the PRIDE database^138^.

### Bioinformatics for plasma proteomics data

Bioinformatic analysis was processed using the Python Jupyter Notebook script (not yet publicly released). PCA, t-SNE, and UMAP were performed to distinguish differences among experimental groups. The three analyses were executed using the scikit-learn library^139^ (v1.5.2) and were visualized using Matplotlib^140^ (v3.9.2). Missing values remaining in the dataset were set to half of the minimum value of each sample. The raw quantifications were normalized using the quantile-normalization method to analyze the difference between samples using EdgeR^141^ (v4.4.2) and Limma^142^ (v3.62.2). Statistical testing was processed using permutation testing (n=10,000). Adjusted *p*-values were computed using two-tailed tests in a combination of t-values, log2 mean difference, and Z-transformed rank-sum statistics, scaling with Liptak Stouffer’s z method^143^. Using the raw data, overall proteome profile similarity was calculated using the Spearman Correlation Coefficient using Scipy^144^ (v1.15.2). Power analysis was performed to assess the statistical power of the study based on the observed sample size and effect size^145^. The median absolute effect size across proteins was used as a representative effect size for power estimation. Statistical power was estimated using a two-sample t-test framework with α = 0.05 and the observed sample sizes and group ratio^146^. Simulation-based power was additionally assessed using 1,000 random simulations based on the observed median effect and pooled standard deviation^147^. The required sample size for 80% power was estimated assuming balanced group sizes.

### Gene properties analysis

GO and KEGG pathway enrichment analysis for the DEPs was performed using DAVID^50^. DEPs were determined with statistical thresholds set at |log_2_FC|>2 and p<0.05. GO and KEGG analyses were conducted using DAVID, with significance defined as an FDR (Benjamini) < 0.05. The top 10 terms with the highest gene counts were selected and plotted.

For SAFE analysis, DEPs were determined with statistical thresholds set at |log_2_FC|>4 and p<0.05. These genes were analyzed using the STRING database (v12.0)^122^, embedded in Cytoscape (v3.10.3)^148^, to construct a protein–protein interaction (PPI) network focused on their predominant GO domains. In STRING, only experimentally supported interactions with a high-confidence score of ≥ 0.7 were included. Visualization of the selected DEPs’ network was marked using Cytoscape (v3.10.3). To visualize this novel network, we used SAFE (v1.0.0) to determine, visualize, and predict significant functional modules in the network^149^. GO terms for each protein were extracted from FuncAssociate_all (v3 -GO updated in February 2018)^150^, and the representative GO term of each cluster was annotated in **Fig. 1D**. The SAFE analysis will be run using Cytoscape with neighborhoodRadius=5, minimum landscape size=10, and landscape Jaccard similarity=0.75.

### Fold-change profile of individual patients

The fold-change of two samples from individual patients was calculated and analyzed by PCA. PCA is applied to determine clinical covariates that contribute to the variability of these samples by the scikit-learn library. The Pearson correlation coefficient was calculated with SciPy to assess the relationship between principal components and clinical covariates. To categorize the DEPs related to sampling interval and LEDD, hierarchical clustering was performed using Euclidean distance after normalization via z-score transformation to ensure a uniform scale across features.

### Data collection from the PPMI database

To investigate more longitudinal proteomics datasets, we downloaded the raw data of proteomics analysis from CSF and Plasma samples obtained at baseline (at the time of diagnosis) and 24-, 36-, 48, 60-, 72-, and 84-month follow-up from the PPMI database^43^. The Patient IDs we analyzed for this study were PP-3123, PP-3372, PP-3377, PP-3378, PP-3752, PP-3814, PP-3819, PP-3825, PP-3826, and PP-3830. The datasets of each patient were separately analyzed in the same way as our plasma proteomics data described above. Then, the fold-change of baseline and each follow-up sample was calculated and analyzed by PCA.

### Prediction tool development using artificial intelligence

To develop the prediction tool, we evaluated a comprehensive panel of supervised machine learning algorithms implemented within the PyTorch framework^151^. The models assessed included three logistic regression models (Ridge, LASSO, ElasticNet)^152,153^. A featureless model was established as a baseline for performance comparison.

### Construction of the validation cohort

We prospectively recruited 38 early-stage PD patients who were diagnosed between January 2017 and January 2021 at the outpatient clinic of the Department of Neurology, Seoul National University Hospital, for validation. 19 Drug-naive patients and 19 drug-exposed patients were enrolled. After informed consent, baseline and follow-up blood samples were collected. To confirm the similarity of discovery and validation cohorts, the Pearson Chi-squared test for sex distribution and the two-sample t-test for other characteristics were performed.

### Sandwich ELISA

A sandwich ELISA was performed for 9 selected candidate markers on samples from the validation cohort, following the manufacturer’s instructions.

| Name | Range (Sensitivity) | Company | Catalog Number |
| --- | --- | --- | --- |
| FN1 | 0.31-20.0 ng/ml (0.1 ng/ml) | Invitrogen | BMS2028 |
| APLP1 | 23.438 - 1500 pg/ml (7.5 pg/ml) | Abcam | ab309112 |
| SRSF3 | 0.156-10ng/ml (0.094ng/ml) | Biorbyt | orb562727 |
| SCAP | 0.156-10ng/ml (0.094ng/ml) | Biorbyt | orb1952971 |
| APAF1 | 0.16-10 ng/mL (0.09 ng/ml) | Bio-Techne | NBP2-68232 |
| EMC4 | 0.156 - 10 ng/ml (<0.05 ng/ml) | Abbexa | abx521961 |
| VWA8 | 78 - 5000 pg/ml (<23.0 pg/ml) | Abbexa | abx553266 |
| DRG1 | 0.312 - 20 ng/ml (<0.14 ng/ml) | Abbexa | abx386986 |
| GPAT4 | 0.312 - 20 ng/ml (< 0.15 ng/ml) | Abbexa | abx385662 |

### PFF uptake screening

The functional genomics screening method followed the protocol from the previous study^28^. The pH-sensitive pHrodo Green conjugated PFF (PFF-pHrodo)^84,85^ was generated as described in previous work ^84^.

We recently established HeLa ORFeome^28^, introducing human ORFeome 9.1^154,155^ and delta space (a total of ∼20,000 ORFs) in a single-gene-per-cell manner using Bxb1 landing pad^156^. The detailed method for establishing an ORFeome-expressing cell line and evaluating the cell line is described in our recent paper^28^.

HeLa ORFeome cells were thawed and pooled to maximize ORF library coverage, then allowed to recover for three days before the experiment. Two million cells, securing 100× coverage, were seeded into each 15 cm cell culture dish in triplicate. On the following day, cells were incubated with 200 ng/ml of PFF-pHrodo for 16 hr at 37 in a CO_2_ incubator. After two DPBS washes, cells were sorted on a BD FACSAria Fusion, gating for mCherry positivity (indicating ORF expression) and then for increased PFF uptake (High green fluorescence, upper ∼10%) and decreased PFF uptake (Low green fluorescence, lower ∼10%) populations. Sorted cells were collected, seeded in 10 cm dishes, and expanded for one week before being cultured to full confluence in 15 cm dishes. Recovered cells were trypsinized and collected (∼10 million cells per pellet) for subsequent genomic DNA extraction. The genomic DNA was extracted using Promega Wizard^®^ Genomic DNA Purification Kit (Promega, A1125), following the manufacturer’s instructions. To amplify ORFs from genomic DNA, we performed two sets of PCRs (nested PCR) using the Q5 High-Fidelity 2X Master Mix (New England Biolabs, M0544S) and primers (Integrated DNA Technologies) that bind to i) the landing pad locus and ii) Gateway-cloning sites, to amplify the full-length ORF from the landing pad with improved selectivity and reduced genomic DNA contamination. Amplified gDNA was quantified using the Quant-iT™ PicoGreen^®^ dsDNA Assay Kit (Thermo Scientific, P7589). Libraries were generated using the NEBNext Ultra II DNA Library Prep Kit for Illumina (New England BioLabs, E7645) as per the manufacturer’s recommendations. Adapters and PCR primers were purchased from Integrated DNA Technologies. The size selection of libraries containing the desired insert size has been performed using SparQ pure beads (Qiagen, 95196-060). Libraries were quantified using the KAPA Library Quantification Kits -Complete kit (Universal) (Kapa Biosystems, KK4824). Average fragment size was determined using a Fragment Analyzer 5300 (Agilent) instrument. The libraries were normalized and pooled and then denatured in 0.02N NaOH and neutralized using pre-load buffer. The pool was loaded at 150 pM on an Illumina NovaSeq X Plus 25B lane following the manufacturer’s recommendations. The run was performed for 2×100 cycles (paired-end mode). A phiX library was used as a control and mixed with libraries at 1% level. Program BCL Convert 4.2.4 was then used to demultiplex samples and generate fastq reads. This ORFeome-seq^28^ was performed by Genome Quebec in Montreal, Canada. The differentially represented ORFs were analyzed by our in-house pipeline^28^. When we defined differentially represented ORFs, the ORFs that were not detected in any group (e.g., detected in the pre-sort group but undetected in the high uptake group) were excluded to avoid false positives.

### Validation of hits with PFF uptake screening

To generate single cell lines for verification, specific entry clones corresponding to ORFs (indicated in Fig. 4B) were grown overnight and isolated using the Qiagen Plasmid Mini Prep kit (Qiagen, 27104) from the ORFeome 9.1 collection^154,155^. 150 ng of entry vector, 150 ng of destination vector (pcDNA, mammalian expression vector), and 1 μl of LR Clonase were incubated for 1 hr and transformed into 10-beta chemically competent cells (New England Biolabs, C3019) using heat shock^157^. Transformants were grown overnight in LB containing carbenicillin (Fisher, 10177012) and used for plasmid extractions. Sequences were confirmed through Sanger reactions. One day before transfection, HeLa cells were seeded into 6-well plates at 300K cells/well. The correctly cloned plasmids were transfected into HeLa cells using Lipofectamine 3000 (Invitrogen, L3000015) according to the manufacturer’s instructions. The media was refreshed after 12 hr to remove the transfection mix. On the following day, 500 ng/ml of PFF-pHrodo were incubated for 16 hr at 37 in a CO_2_ incubator. After two washes with DPBS, cells were analyzed using a BD LSRFortessa X-20 to measure green pHrodo fluorescence as an indicator of PFF uptake. The pHrodo-positive cell population was normalized by the empty vector control. The graph was generated using GraphPad Prism 9, and A two-tailed t-test was performed to assess the statistical significance.

### PFF-induced toxicity screening with stem cell-derived dopamine-positive neurons

#### Cell lines and culture conditions

H9 embryonic stem cell lines were maintained in Stemflex medium (Stemflex Medium, Catalogue number: Gibco A3349401) on Matrigel (Corning^®^ Matrigel^®^ hESC-Qualified Matrix, Catalogue number: 354277) coated dishes. Cells were replated every 4-5 days with 0.5 mM EDTA (Invitrogen, AM9260G).

#### Lentivirus transduction

Confluent H9 cells were dissociated into single cells with Accutase (Stemcell Tech, 07922) and plated on Matrigel-coated dishes at a density of 1.5 million cells per well in a 6-well plate in StemFlex medium with ROCK inhibitor Y27632 (Selleckchem, s1049). Cells were left to settle for 2 hr and then treated with lentivirus with 1000X polybrene (Sigma-Aldrich, TR-1003-G). After 3 h, cells were washed 2 times with DMEM/F12 (Thermo Fisher, 11330057), and the media was replaced with fresh Stemflex with ROCK inhibitor (Selleckchem, Y-27632).

#### Dopamine neuron differentiation

The differentiation protocol was adapted from a previous protocol^158^. 20 μl of Matrigel (Corning^®^ Matrigel^®^ hESC-Qualified Matrix; Corning, 354277) was plated on 6 spots on a 6 cm plate and left to incubate for 30 min to 1 hr at 37 °C. H9 embryonic stem cells were dissociated with Accutase (Stemcell Tech, 07922) to single cells, and the concentration of cells was diluted with Stemflex (Stemflex Medium; Gibco, A3349401) medium supplemented with ROCK inhibitor (Selleckchem, s1049) to 1 million cells per milliliter. Matrigel was removed from the spots, and 20 μl of this cell suspension was added to the Matrigel-coated spots. The cells were left to settle for 20 min at 37 °C, after which 5 ml of Stemflex supplemented with ROCK inhibitor was added to the plate. The next day (Day 1), the media was fully changed to Day 1 media.

From day 1 to 6, cells were grown in floor plate induction medium in DMEM media (Gibco, LM 001-05) with 15% KOSR (Gibco, 10828-028), Glutamax (Gibco, 25050079), and β-mercaptoethanol (Gibco, 21985023). From day 6 onwards, cells were grown in neural precursor induction medium in DMEM with Glutamax, NEAA, beta mercapto-ethanol, and the following: 11.5% KOSR and 0.25% N2 (Gibco, 17502048) (Day 6-8), 7.5% KOSR and 0.5% N2 (Day 8-10), 3.75% KOSR and 0.75% N2 (Day 10-12). Dual Smad inhibitors 0.2 μM LDN193189 (Stemolecule, 04-0074) and 10 μM SB431542 (Stemcelltech, 72234) were added from Days 1-12, and Days 1-8, respectively. SHH agonists 2 μM Purmorphamine (Sigma-Aldrich, SML0868-5MG) and 100 ng/ml Shh (Peprotech, 100-45-1MG), along with 100 ng/ml FGF8 (Peprotech, 100-25-100UG), were added from Days 2-10. The Wnt signaling activator 1 μM CHIR99021 (Stemcelltech, 72054) was added from Day 4-12.

From day 12 onwards, cells were grown in Dopamine progenitor induction and maturation medium with DMEM:F12 medium with N2, 20 ng/ml BDNF (Peprotech, 450-02), 20 ng/ml GDNF (Peprotech, 450-10), 500 μM dbcAMP (Selleckchem, S7858), 200 μM ascorbic acid (Sigma-Aldrich, A4544), and 10 ng/ml TGF-3 (Peprotech, 100-36E). 10 μM DAPT (Caymen Chem, 13197) and 1 μM CHIR99021 (Stemcelltech, ST72052) were included from Day 12-15. Spots were dissociated on Day 15 and replated in Poly-L-ornithine/Fibronectin/Laminin-coated (PLO: Sigma-Aldrich, p4957; Fibronectin: Sigma Aldrich, F0895; Laminin: Sigma-Aldrich, L2020) dishes at 5 million cells per 60 phi dish. 10 μM Y-27632 (ROCK inhibitor) was added to the medium on dissociation day 15.

#### PFF-induced toxicity Screening

Differentiated cells transfected with lentivirus were treated with doxycycline (StemCellTech, 72742) at 2 μg/ml starting from day 18. Cells were exposed to in vitro cell stress conditions of 5 μg/ml of PFF for 1 week before sampling. PFF was prepared according to a previously published protocol^159^. Genomic DNA was extracted using the Qiawave DNA Blood and Tissue Kit (Qiagen, 69554), and ORFs were PCR-amplified with primers specific to the lentiviral vector with Q5 High-Fidelity 2x polymerase (NEB, M0492L). Amplified DNA was quantified using Qubit dsDNA HS Assay kit (Q32854), and submitted to Genome Québec (Montreal, Canada) for library preparation and sequencing. Libraries were sequenced on an Illumina NovaSeq instrument using paired-end 100 bp reads with a target depth of approximately 25 million reads per sample. Demultiplexed FASTQ files were used for downstream ORFeome-seq analysis as described above.

#### Validation of hits with PFF-induced toxicity screening

H9 human embryonic stem cells (hESCs) were transduced with lentiviruses expressing single ORFs that had been LR-cloned (Invitrogen, 11791100) into a dox-inducible vector. Transduced cells underwent selection using 0.5 μg/ml puromycin (Sigma Aldrich, P9620-10ML) for one week before differentiation into dopaminergic neurons, as previously described^158^. On day 21 of differentiation, the neurons were treated with either 5 μg/ml of PFFs, which were probe-sonicated immediately before use, or a DPBS vehicle control. Cells were subsequently harvested for cytotoxicity assays 14 days post-exposure.

#### Immunofluorescence labeling and quantification

For immunocytochemical analysis, both PFF-and vehicle-treated dopaminergic neurons were fixed in 4% PFA (ISS, SM-P01-100). The cells were then blocked and permeabilized for 1 hr at room temperature (RT) in a buffer containing 5% donkey serum (Merck, S30-100M) and 0.1% Triton X-100 (Sigma Aldrich, X100). Following blocking, neurons were incubated overnight at 4°C with a primary anti-tyrosine hydroxylase (TH) polyclonal antibody (Sigma Aldrich, AB1542). Cells were subsequently incubated with fluorescent Alexa Fluor secondary antibodies (Jackson ImmunoResearch, 713-545-003) for 2 hr at RT and counterstained with Hoechst (Invitrogen, H3570) before mounting. Fluorescence imaging was performed at 20x magnification using a Nikon A1 microscope. Finally, dopaminergic neuron survival was quantified using ImageJ software to count the TH-positive and Hoechst-positive cells.

### Genetic cross-referencing

We compared our proteomics data and functional screening results with PD-associated CNVs reported in recent literature^106^. The intersection between these CNVs and our experimentally identified hits was assessed using Fisher’s exact test, implemented by MATLAB R2018a. Additionally, we compared our proteomics data and snRNA-seq results from the GEO database (GSE243639^107^ and GSE193688^108^). The DEGs in dopaminergic neurons were compared with our DEP set to identify overlapping candidates.

### Meta-analysis of the dopamine treatment effect on *in vivo* mouse brain models

For meta-analysis, we gathered differentially expressed genes of each of the three public datasets from the GEO database (GSE291024^111^ and GSE279704) and Zenodo (#14762864)^112^. The DEGs were compared with our LEDD-dependent DEP set to identify overlapping candidates.

### Network analysis of differentially expressed proteins from proteomics and candidates from functional screening of PFF uptake and cytotoxicity

To perform integrative network analysis using SAFE^149^, we aggregated candidates from our three primary datasets. First, plasma proteins discriminative for the sampling interval were selected based on an absolute difference between subgroups of >1 and p<0.05. Second, functional regulators from the PFF uptake screen were identified using thresholds of |log_2_FC|>0.58 and p<0.05. Last, functional regulators from the PFF-induced neuronal toxicity screen were identified using thresholds of |log_2_FC|>0.32 and p<0.05. These candidates were mapped onto a PPI network using the STRING database^122^, incorporating all active interactions with a medium confidence score of ≥ 0.4. The network topology was generated using an edge-weighted spring-embedded layout, and only the largest connected component was retained for functional annotation via SAFE, following the methods described above. The subnetwork of the proteins exhibiting significant changes in at least two datasets with a maximum neighborhood score was generated using the STRING database, incorporating all active interactions with a low confidence score of ≥ 0.15.

